# Linking Pregnancy- and Birth-Related Risk Factors to a Multivariate Fusion of Child Cortical Structure

**DOI:** 10.1101/2024.10.29.620834

**Authors:** Linn R. S. Lindseth, Dani Beck, Lars T. Westlye, Christian K. Tamnes, Linn B. Norbom

**Affiliations:** PROMENTA Research Center, Department of Psychology, University of Oslo, Oslo, Norway; Division of Mental Health and Substance Abuse, Diakonhjemmet Hospital, Oslo, Norway; Department of Psychology, University of Oslo, Oslo, Norway; Center for Precision Psychiatry, Division of Mental Health and Addiction, Oslo University Hospital, Oslo, Norway; KG Jebsen Centre for Neurodevelopmental Disorders, University of Oslo, Oslo, Norway

**Keywords:** Cortical morphology, MRI, Neurodevelopment, Prenatal, Perinatal

## Abstract

Pregnancy- and birth-related factors affect offspring brain development, emphasizing the importance of early life exposures. While most previous studies have focused on a few variables in isolation, here we investigated associations between a broad range of pregnancy- and birth-related variables and multivariate cortical brain MRI features. Our sample consisted of 8,396 children aged 8.9 to 11.1 years from the ABCD Study. Through multiple correspondence analysis and factor analysis of mixed data, we distilled numerous pregnancy and birth variables into four overarching dimensions; maternal pregnancy complications, maternal substance use, low birth weight and prematurity, and newborn birth complications. Vertex-wise measures of cortical thickness, surface area, and curvature were fused using linked independent component analysis. Linear mixed effects models showed that maternal pregnancy complications and low birth weight and prematurity were associated with smaller global surface area. Additionally, low birth weight and prematurity was associated with complex regional cortical patterns reflecting bidirectional variations in both surface area and cortical thickness. Newborn birth complications showed multivariate patterns reflecting smaller occipital- and larger temporal area, bi-directional frontal area variations, and reduced cortical thickness across the cortex. Maternal substance use showed no associations with child cortical structure. By employing a multifactorial and multivariate morphometric fusion approach, we connected complications during pregnancy and fetal size and prematurity to global surface area and specific regional signatures across child cortical MRI features.

**Significance Statement:** Early life stages, including pregnancy and birth circumstances, are critically important for a wide range of real-life outcomes for the child. In this study, we linked maternal complications during pregnancy and low birth weight and prematurity to globally reduced cortical surface area, and low birth weight and prematurity, in addition to pregnancy and birth complications, to complex cortical structural patterns later in late childhood. Our findings underscore the importance of providing support to mothers and children during these crucial phases, helping to ensure optimal conditions for healthy child development.

## Introduction

Decades of research document that the period from conception to birth is crucial for the offspring’s subsequent development (1, 2). Adverse events and exposures during this time have been linked to a variety of negative outcomes, including lower general cognitive ability (3–6), behavioral problems (6–10), and neurodevelopmental conditions (11, 12).

During pregnancy the cerebral cortex of the fetus undergoes rapid maturation (13), and characteristics of cortical structure at birth have been shown to predict later function (14). The cortical plate begins forming around 7 weeks post-conception (15, 16), and by 20 weeks of gestation it can be observed as a smooth surface in fetal magnetic resonance imaging (MRI) (17). The greatest increase in cortical thickness (CT) occurs mid-gestation, reaching about 80% of its maximum size by birth, while surface area (SA) expands from mid-gestation, reaching approximately 25% of its maximum size at birth (13). Most of the cortical folding occurs during the final trimester of pregnancy (17, 18).

Postnatal development of the cerebral cortex is characterized by a continued thickening for the first two years of life (13, 19, 20), followed by a gradual thinning throughout childhood and adolescence (13, 21, 22). SA increases rapidly the first two years of life (13, 23) and continues to expand at a slower pace until middle childhood (13), followed by a subtle decrease through adolescence (13, 21, 22). Increases in cortical folding during the first two years of life (24) are followed by slow decreases (21). Cortical development also exhibits both individual (25, 26) and sex (21) differences, and studies have shown that cortical morphology is highly heritable (27). Despite this strong genetic influence, environmental factors and experiences can fundamentally shape cortical structure.

A wide range of early-life environmental risk factors have been linked to variations in child cortical morphology in childhood and adolescence, though findings have been inconsistent. Maternal pregnancy-related complications such as pre-eclampsia and high blood-pressure have been associated with both increased (9, 28) and decreased (3) CT, as well as smaller SA (6, 28). Prenatal exposure to substances has been associated with bot increased (10, 29–32) and decreased (7, 10, 31) CT, and both larger (8) and smaller (29, 33, 34) SA. Further, a range of birth-related variables, including caesarian delivery, being born before due date, and lower birth weight, have been related to increased CT (28, 35–39), though lower birth weight has also been associated with decreased CT (5, 37–39). Several birth-related variables have also been associated with smaller SA (5, 28, 35–40). Although research on cortical folding is more limited, studies suggest that various pregnancy- and birth-related variables are associated with altered sulcal patterns (10, 28) and gyrification (36).

Our current understanding of the complex relationships between pregnancy- and birth-related variables and future cortical structure in the offspring is limited by the fact that most previous studies have focused on one or a few variables and one or a few brain metrics in isolation. In contrast, investigating a broad set of variables related to pregnancy and birth circumstances could provide insight into the potentially overlapping effects of diverse risk factors. Further, it is likely that multiple biological processes, here captured by partly separate cortical metrics, are both uniquely and collectively influenced by pregnancy- and birth-related factors. Indeed, a recent study that assessed a wide range of environmental variables along with multiple neuroimaging metrics revealed complex relationships between pregnancy- and birth-related factors and child brain structure (28).

By examining numerous risk factors across pregnancy and birth in relation to multiple cortical metrics, our aim was to disentangle heterogeneous findings in previous research and provide a more comprehensive understanding of the influence of these factors on child brain structure. In a sample of 8,396 participants aged 8.9 to 11.1 years (Mean = 9.92, SD = 0.63, 47.9% female) from the Adolescent Brain Cognitive Development (ABCD) Study (https://abcdstudy.org/), we tested the associations between a broad set of pregnancy- and birth-related risk factors and child cortical structure, assessed through fusion of CT, SA, and curvature. Fusion was performed using a linked independent component analysis (Linked-ICA) (41) approach, which decomposes cortical surface maps into statistically independent components, capturing both shared (multivariate) and unique (univariate) patterns of variation, providing a data-driven representation of structural organization. Pregnancy and birth variables included maternal somatic health in pregnancy, substance use, and birth complications. We hypothesized that exposures to pregnancy- and birth-related risk factors would be linked to higher CT (9, 28–32) and lower SA (6, 28, 33–35, 37–40, 42) of the child, grossly reflecting a less mature brain pattern in late childhood. Additionally, we expected pregnancy- and birth-related risk to be associated with regionally altered cortical curvature (10, 28, 36).

## Results

### Pregnancy- and Birth-Related Dimensions

Using the ABCD Developmental History Questionnaire (43), we identified 23 pregnancy and 12 birth variables, which were reduced to overarching dimensions using Factor Analysis of Mixed Data (FAMD) and Multiple Correspondence Analysis (MCA). For further analyses, we selected the first two dimensions from both reduction methods, which explained 9.2% and 7.1% of the total pregnancy-related variance, and 22.8% and 14.4% of the total birth-related variance, respectively (SI Appendix Fig. S1). We labeled the first pregnancy dimension “Maternal Pregnancy Complications” as it comprised variables relating to maternal somatic health during pregnancy, including occurrence of rubella, proteinuria, high blood-pressure, and (pre-)eclampsia. We labeled the second pregnancy dimension “Maternal Substance Use”, which included variables related to use of tobacco, marijuana, and alcohol, in addition to other substances. We named the first birth dimension “Low Birth Weight and Prematurity” as the most contributing variables were weeks born before due date and birth weight. The second birth dimension, “Newborn Birth Complications”, was characterized by variables such as the child appearing blue in color, not breathing, or having a slow heartbeat. The most contributing variables and their explained variance in the given dimensions are shown in Fig. 1, while density plots for the selected dimensions are shown in SI Appendix Fig. S2. Further details on the results of the reduction analysis are presented in the “Methods” section of the SI Appendix. Loadings of the contributing variables in the dimensional space are illustrated in in SI Appendix Fig. S3.

**Figure 1.**
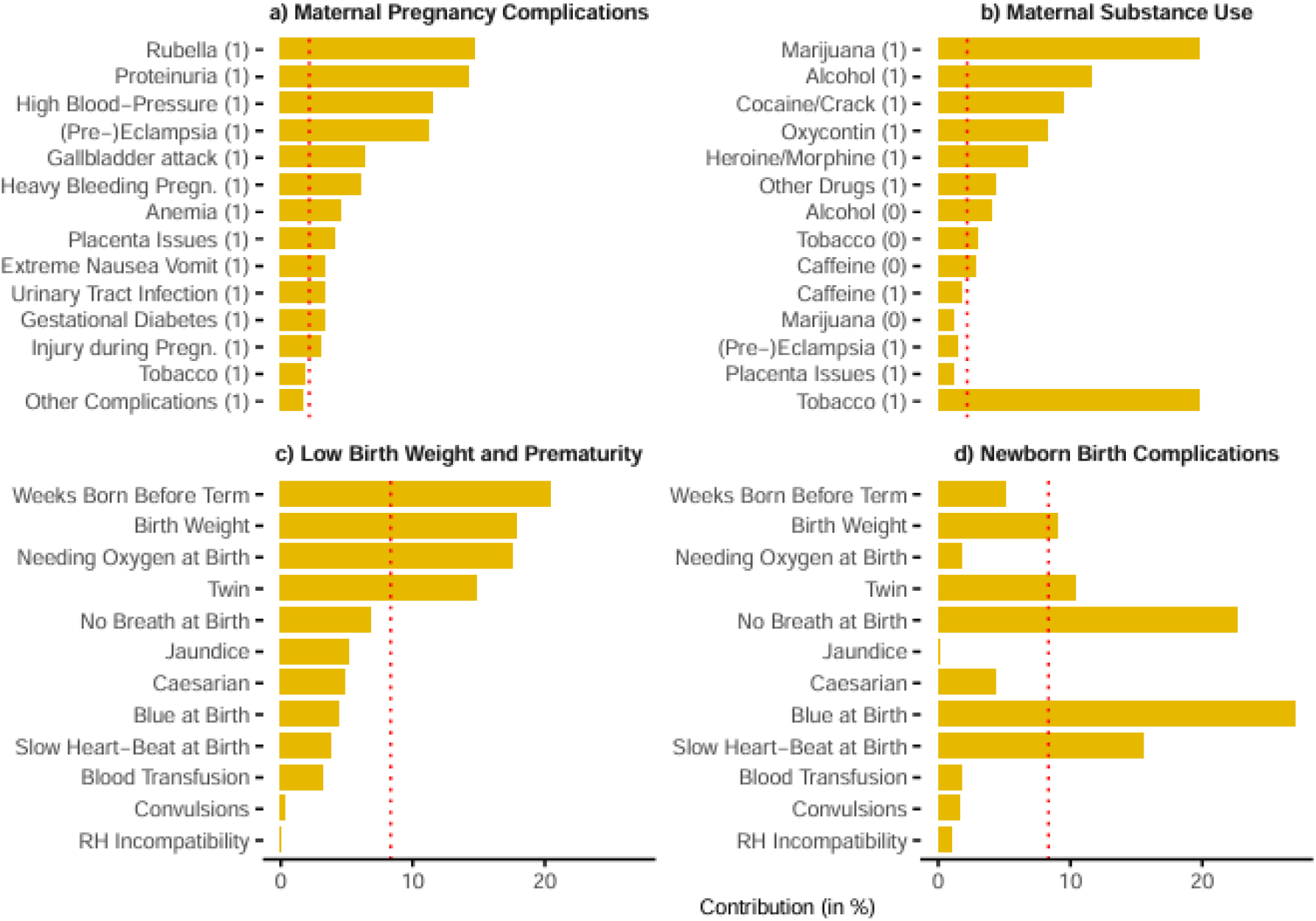
Variable contribution to pregnancy- and birth-related dimensions. The figure shows the variables with the highest contributions to each dimension. The x-axis indicates the percentage contribution while the red dotted line represents the expected average contribution if the variables were uniform. a) and b) are pregnancy dimensions while c) and d) are birth dimensions. (1) and (0) indicates whether the response to the variable is yes or no, respectively. Figure a) and b) show the top 14 contributing variables in the given dimension.

### Cortical Decomposition

Final decomposition of vertex-wise surface maps of CT, SA, and curvature yielded 39 independent cortical components (CC) (Fig. 2). In general, SA contributed most to the components overall, followed by CT, while curvature only had minor contributions. The components captured both unique and multivariate patterns, and the respective weighting and explained variance are presented in SI Appendix Tab. S1, while surface maps for all CCs are presented in SI Appendix Fig. S4-7. CC1 accounted for 29.30% of the total variance in the imaging data, while the remaining CCs explained between 7.26% and 0.99%.

**Figure 2.**
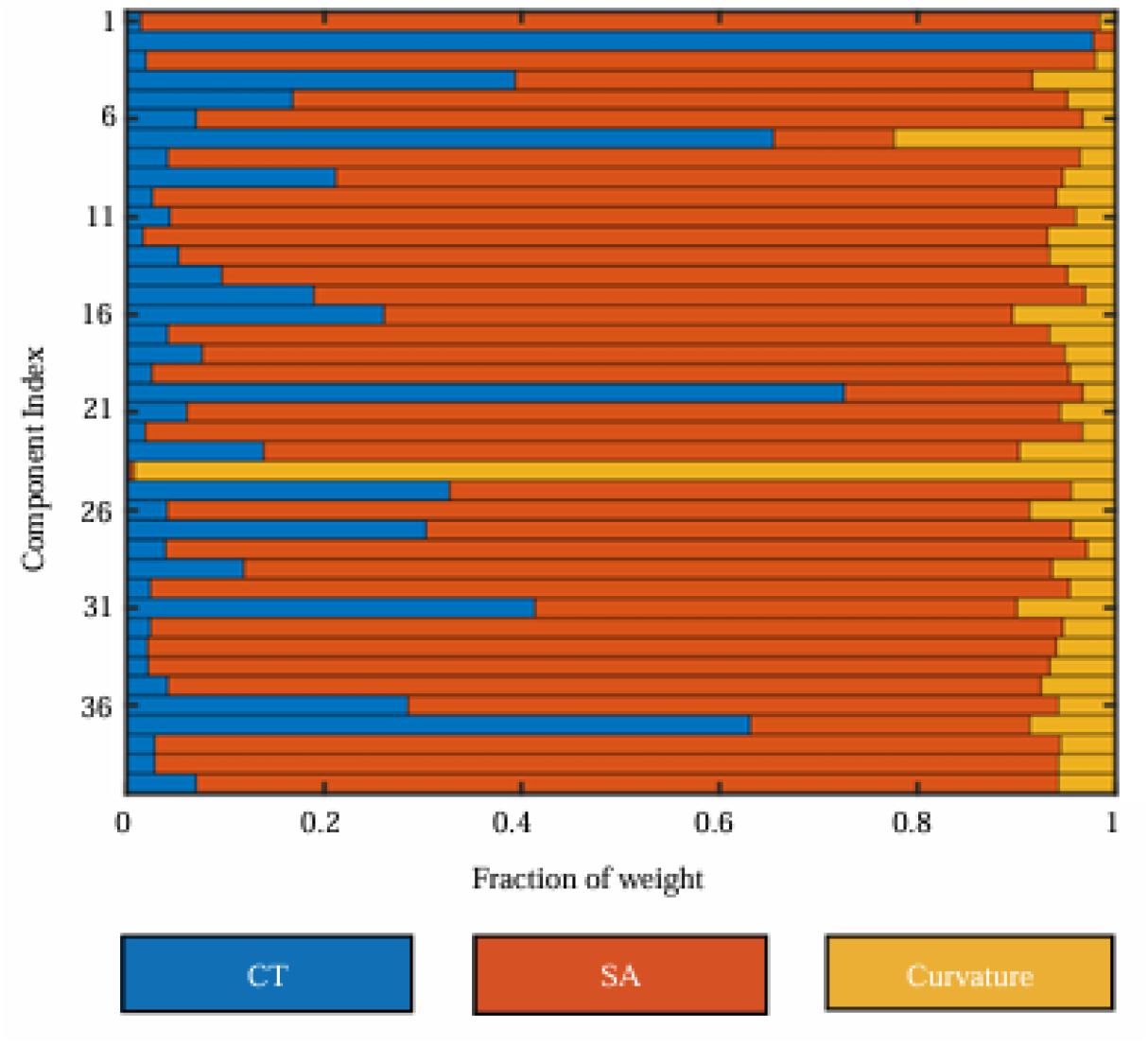
Cortical fusion decomposition. The figure shows the decomposition of the 40 cortical components, with relative weight of each of the three brain metrics CT (cortical thickness), SA (surface area), and Curvature.

### Associations Between Pregnancy-Related Dimensions and Cortical Structure

Linear mixed effects modelling revealed a significant negative association between Maternal Pregnancy Complications and CC1 (t = −3.81, Cohen’s D = −0.09, p_adjusted_ < 0.001) (Tab. 1 and Fig. 3). CC1 was dominated by global SA (97%), indicating that mothers who experienced somatic health complications during pregnancy had children with smaller global SA at approximately age 10 compared to mothers with fewer such complications.

**Table 1.** Associations between pregnancy-related dimensions and CCs.

| CC | Maternal Pregnancy Complications |  |  |  | Maternal Substance Use |  |  |  |
| --- | --- | --- | --- | --- | --- | --- | --- | --- |
|  | Cohen's D | t-value | Non-parametric p-value (unadj.) | Non-parametric p-value (adj.) | Cohen's D | t-value | Non-parametric p-value (unadj.) | Non-parametric p-value (adj.) |
| 1 | -0.09 | -3.81 | **<0.001** | **<0.001** | 0.02 | 0.78 | 0.430 | 0.959 |
| 2 | -0.05 | -2.30 | **0.021** | 0.763 | 0.02 | 0.94 | 0.332 | 0.959 |
| 3 | -0.02 | -0.88 | 0.378 | 0.864 | 0.03 | 1.42 | 0.146 | 0.959 |
| 4 | 0.02 | 0.68 | 0.493 | 0.864 | -0.04 | -1.71 | 0.087 | 0.959 |
| 5 | -0.01 | -0.56 | 0.570 | 0.864 | 0.02 | 0.84 | 0.398 | 0.959 |
| 6 | 0.02 | 0.88 | 0.373 | 0.864 | 0.03 | 1.12 | 0.262 | 0.959 |
| 7 | -0.02 | -0.90 | 0.377 | 0.864 | 0.00 | -0.20 | 0.833 | 0.959 |
| 8 | 0.01 | 0.30 | 0.771 | 0.864 | -0.02 | -0.85 | 0.400 | 0.959 |
| 9 | -0.02 | -0.69 | 0.485 | 0.864 | -0.05 | -2.30 | **0.022** | 0.851 |
| 10 | -0.05 | -2.13 | **0.034** | 0.864 | 0.02 | 1.04 | 0.303 | 0.959 |
| 11 | -0.01 | -0.29 | 0.770 | 0.864 | 0.00 | 0.14 | 0.893 | 0.959 |
| 12 | 0.03 | 1.34 | 0.180 | 0.864 | 0.00 | -0.05 | 0.959 | 0.959 |
| 13 | -0.02 | -0.82 | 0.413 | 0.864 | -0.02 | -1.02 | 0.303 | 0.959 |
| 14 | -0.02 | -0.86 | 0.386 | 0.864 | 0.02 | 1.01 | 0.309 | 0.959 |
| 15 | -0.04 | -1.74 | 0.084 | 0.864 | -0.01 | -0.49 | 0.619 | 0.959 |
| 16 | 0.02 | 1.03 | 0.296 | 0.864 | -0.04 | -1.54 | 0.121 | 0.959 |
| 17 | -0.01 | -0.59 | 0.563 | 0.864 | -0.01 | -0.44 | 0.659 | 0.959 |
| 18 | 0.00 | -0.20 | 0.847 | 0.864 | 0.00 | 0.17 | 0.863 | 0.959 |
| 19 | -0.05 | -2.15 | **0.037** | 0.864 | -0.02 | -0.92 | 0.351 | 0.959 |
| 20 | -0.06 | -2.58 | **0.012** | 0.459 | -0.01 | -0.29 | 0.773 | 0.959 |
| 21 | -0.02 | -0.91 | 0.354 | 0.864 | 0.04 | 1.61 | 0.107 | 0.959 |
| 22 | 0.02 | 0.65 | 0.518 | 0.864 | -0.05 | -2.26 | **0.026** | 0.955 |
| 23 | 0.01 | 0.36 | 0.727 | 0.864 | 0.00 | 0.13 | 0.908 | 0.959 |
| 25 | 0.00 | 0.17 | 0.864 | 0.864 | 0.03 | 1.17 | 0.241 | 0.959 |
| 26 | -0.01 | -0.27 | 0.787 | 0.864 | 0.00 | 0.17 | 0.877 | 0.959 |
| 27 | -0.05 | -1.92 | 0.058 | 0.864 | 0.02 | 0.87 | 0.375 | 0.959 |
| 28 | -0.02 | -0.75 | 0.454 | 0.864 | -0.04 | -1.65 | 0.101 | 0.959 |
| 29 | -0.04 | -1.65 | 0.099 | 0.864 | -0.04 | -1.81 | 0.072 | 0.959 |
| 30 | 0.01 | 0.29 | 0.768 | 0.864 | -0.03 | -1.51 | 0.143 | 0.959 |
| 31 | -0.01 | -0.50 | 0.614 | 0.864 | -0.05 | -2.03 | **0.044** | 0.959 |
| 32 | -0.05 | -2.22 | **0.025** | 0.864 | 0.01 | 0.42 | 0.674 | 0.959 |
| 33 | 0.05 | 2.13 | **0.034** | 0.864 | 0.03 | 1.35 | 0.170 | 0.959 |
| 34 | -0.01 | -0.35 | 0.734 | 0.864 | 0.05 | 2.26 | **0.027** | 0.959 |
| 35 | 0.07 | 2.88 | **0.003** | 0.122 | 0.02 | 1.00 | 0.322 | 0.959 |
| 36 | -0.02 | -0.93 | 0.349 | 0.864 | 0.01 | 0.55 | 0.569 | 0.959 |
| 37 | 0.02 | 0.77 | 0.430 | 0.864 | 0.06 | 2.65 | **0.010** | 0.398 |
| 38 | -0.01 | -0.25 | 0.799 | 0.864 | -0.01 | -0.25 | 0.792 | 0.959 |
| 39 | -0.01 | -0.41 | 0.688 | 0.864 | 0.01 | 0.32 | 0.744 | 0.959 |
| 40 | 0.01 | 0.35 | 0.725 | 0.864 | -0.03 | -1.50 | 0.136 | 0.959 |

The table shows effect sizes, t-values, and the non-parametric uncorrected (Unadj.) and corrected (Adj.) p-values. Significant p-values with threshold 0.05 are shown in bold.

**Figure 3.**
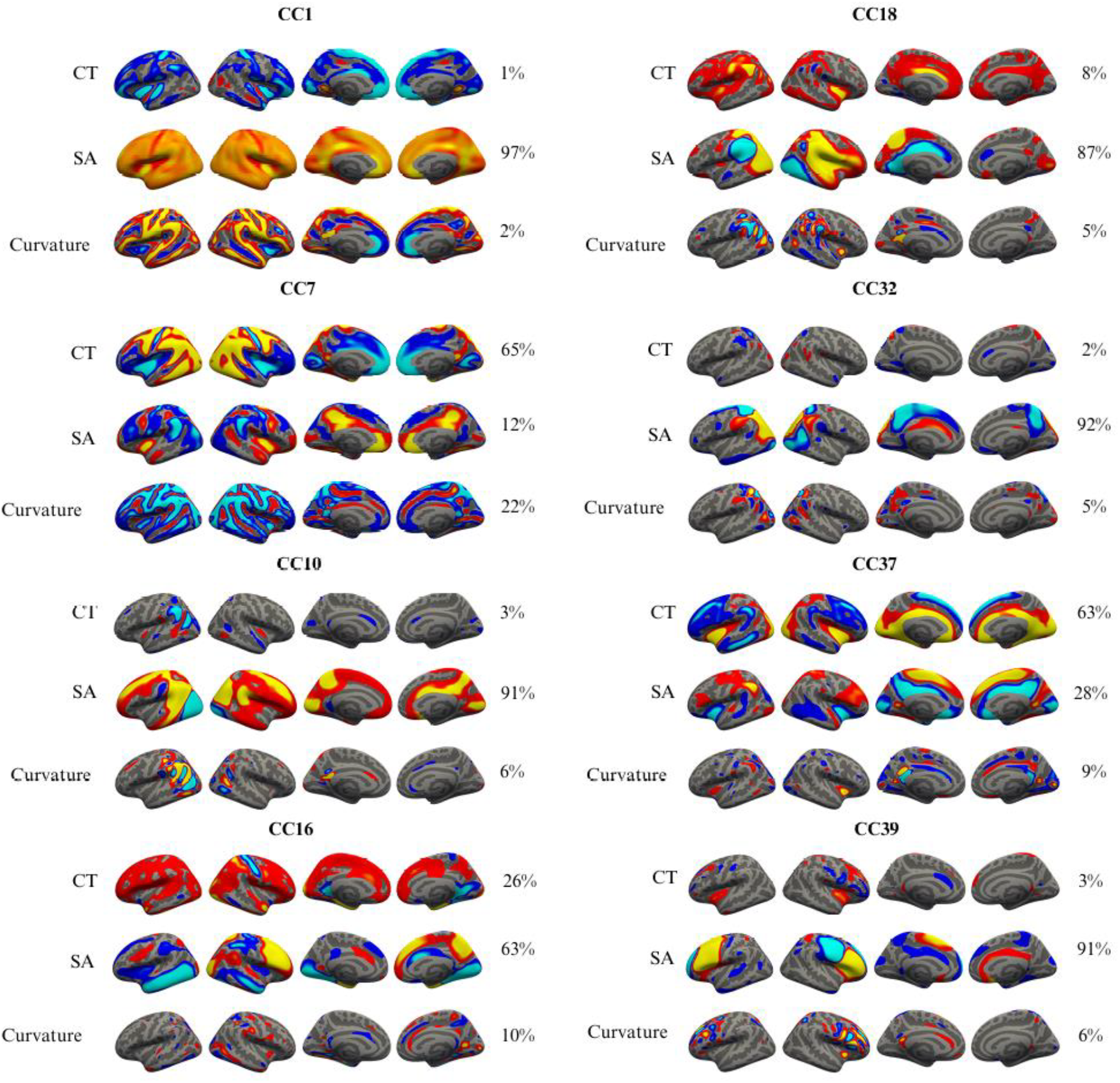
Cortical components associated with pregnancy- and birth-related dimensions. The figure shows the cortical components (CC)s with a significant association with pregnancy- and/or birth-related dimensions. All surface maps are thresholded with a minimum of 4 and maximum of 15 standard deviations, except for surface area within CC1, which were thresholded with a minimum of 20 and a maximum of 100, to enhance regional detail within the global pattern. Percentages represent the contributions of each metric within the CC.

There were no significant associations between either Maternal Substance Use and any of the CCs (Tab. 1).

### Associations Between Birth-Related Dimensions and Cortical Structure Low Birth Weight and Prematurity

Linear mixed effects modelling revealed significant associations between Low Birth Weight and Prematurity and CC1, CC7, CC16, CC18, CC32, CC37 and CC39 (Tab. 2 and Fig. 3).

**Table 2.** Associations between birth-related dimensions and CC.

| CC | Low Birth Weight and Prematurity |  |  |  | Newborn Birth Complications |  |  |  |
| --- | --- | --- | --- | --- | --- | --- | --- | --- |
|  | Cohen's D | t-value | p-value (unadj.) | p-value (adj.) | Cohen's D | t-value | p-value (unadj.) | p-value (adj.) |
| 1 | -0.14 | -5.86 | **<0.001** | **<0.001** | 0.06 | 2.57 | **0.009** | 0.308 |
| 2 | -0.04 | -1.65 | 0.099 | 0.910 | -0.01 | -0.52 | 0.599 | 0.993 |
| 3 | 0.01 | 0.24 | 0.820 | 0.910 | -0.03 | -1.25 | 0.210 | 0.993 |
| 4 | -0.07 | -2.86 | **0.003** | 0.105 | 0.03 | 1.41 | 0.163 | 0.993 |
| 5 | -0.03 | -1.33 | 0.185 | 0.910 | 0.01 | 0.38 | 0.705 | 0.993 |
| 6 | -0.02 | -0.58 | 0.560 | 0.910 | 0.04 | 1.93 | 0.055 | 0.993 |
| 7 | -0.09 | -3.44 | **<0.001** | **0.014** | 0.05 | 2.37 | **0.018** | 0.576 |
| 8 | 0.01 | 0.44 | 0.656 | 0.910 | -0.03 | -1.56 | 0.130 | 0.993 |
| 9 | 0.07 | 2.95 | **0.003** | 0.090 | -0.01 | -0.59 | 0.558 | 0.993 |
| 10 | -0.06 | -2.37 | **0.021** | 0.615 | 0.08 | 3.52 | **0.001** | **0.023** |
| 11 | 0.00 | -0.15 | 0.888 | 0.910 | 0.00 | 0.01 | 0.993 | 0.993 |
| 12 | 0.00 | 0.11 | 0.910 | 0.910 | 0.01 | 0.24 | 0.807 | 0.993 |
| 13 | 0.03 | 1.40 | 0.165 | 0.910 | -0.01 | -0.29 | 0.776 | 0.993 |
| 14 | -0.01 | -0.25 | 0.808 | 0.910 | 0.01 | 0.37 | 0.716 | 0.993 |
| 15 | -0.06 | -2.48 | **0.012** | 0.372 | 0.04 | 1.79 | 0.081 | 0.993 |
| 16 | 0.16 | 6.17 | **<0.001** | **<0.001** | -0.10 | -4.41 | **<0.001** | **<0.001** |
| 17 | 0.01 | 0.40 | 0.683 | 0.910 | -0.02 | -0.96 | 0.326 | 0.993 |
| 18 | -0.09 | -3.42 | **0.001** | **0.020** | 0.07 | 3.19 | **0.002** | 0.059 |
| 19 | -0.02 | -0.83 | 0.412 | 0.910 | -0.06 | -2.58 | **0.011** | 0.367 |
| 20 | -0.02 | -0.60 | 0.543 | 0.910 | 0.00 | -0.07 | 0.946 | 0.993 |
| 21 | -0.03 | -1.29 | 0.196 | 0.910 | 0.03 | 1.18 | 0.248 | 0.993 |
| 22 | 0.03 | 1.23 | 0.213 | 0.910 | -0.01 | -0.43 | 0.661 | 0.993 |
| 23 | -0.03 | -1.42 | 0.156 | 0.910 | -0.01 | -0.59 | 0.546 | 0.993 |
| 25 | 0.02 | 0.86 | 0.389 | 0.910 | -0.01 | -0.62 | 0.530 | 0.993 |
| 26 | -0.05 | -1.82 | 0.066 | 0.910 | 0.03 | 1.55 | 0.126 | 0.993 |
| 27 | -0.04 | -1.35 | 0.192 | 0.910 | 0.04 | 1.62 | 0.106 | 0.993 |
| 28 | -0.01 | -0.47 | 0.628 | 0.910 | 0.04 | 2.00 | **0.048** | 0.993 |
| 29 | 0.02 | 0.70 | 0.495 | 0.910 | -0.01 | -0.48 | 0.636 | 0.993 |
| 30 | -0.02 | -0.67 | 0.501 | 0.910 | -0.01 | -0.32 | 0.753 | 0.993 |
| 31 | 0.03 | 1.25 | 0.220 | 0.910 | -0.05 | -2.14 | **0.034** | 0.993 |
| 32 | -0.12 | -4.60 | **<0.001** | **<0.001** | 0.05 | 2.29 | **0.020** | 0.614 |
| 33 | 0.05 | 2.04 | **0.042** | 0.910 | -0.06 | -2.82 | **0.005** | 0.166 |
| 34 | -0.06 | -2.24 | **0.025** | 0.706 | 0.02 | 0.76 | 0.432 | 0.993 |
| 35 | 0.03 | 1.24 | 0.214 | 0.910 | -0.03 | -1.58 | 0.119 | 0.993 |
| 36 | 0.04 | 1.77 | 0.072 | 0.910 | -0.03 | -1.41 | 0.156 | 0.993 |
| 37 | 0.14 | 5.71 | **<0.001** | **<0.001** | -0.03 | -1.23 | 0.212 | 0.993 |
| 38 | 0.02 | 0.89 | 0.364 | 0.910 | -0.03 | -1.40 | 0.162 | 0.993 |
| 39 | -0.13 | -5.23 | **<0.001** | **<0.001** | 0.05 | 2.43 | **0.015** | 0.488 |
| 40 | 0.01 | 0.51 | 0.590 | 0.910 | -0.02 | -0.95 | 0.338 | 0.993 |

The table shows effect sizes, t-values, and the non-parametric uncorrected (Unadj.) and corrected (Adj.) p-values. Significant p-values with threshold 0.05 are shown in bold.

Low Birth Weight and Prematurity was negatively associated with CC1 (t = −5.8, Cohen’s D = −0.14, p_adjusted_ < 0.001), indicating that children with lower birth weight and/or prematurity had smaller global SA at around 10 years of age.

Low Birth Weight and Prematurity showed a negative association with CC7 (t = −3.44, Cohen’s D = −0.09, p_adjusted_ = 0.014). CC7 reflected a multivariate pattern of CT (65%), curvature (22%) and SA (12%). The results indicate that beyond the global findings already described, children with lower birth weight and/or being born premature have reduced parietal, occipital and temporal CT, thicker and smaller insular cortex, and mainly higher, with some regions of lower, curvature spread across the cortex in late childhood.

Low Birth Weight and Prematurity showed a positive association with CC16 (t = 6.17, Cohen’s D = 0.16, p_adjusted_ < 0.001). CC16 reflected a multivariate pattern of SA (63%), CT (26%) and curvature (10%). These results imply that children with lower birth weight and/or being born premature have somewhat smaller temporal and insular SA, larger right-hemispheric frontal SA, and a pattern of increased CT spread across cortical regions.

Low Birth Weight and Prematurity showed a negative association with CC18 (t = −3.42, Cohen’s D = −0.09, p_adjusted_ = 0.020). CC18 reflected a more univariate pattern of SA (87%) with smaller contributions from CT (8%) and curvature (5%). These results show that children with lower birth weight and/or being born premature have smaller left-hemispheric and larger right-hemispheric occipital SA, and somewhat reduced insular, frontal and parietal CT.

Low Birth Weight and Prematurity showed a negative association with CC32 (t = −4.60, Cohen’s D = −0.12, p_adjusted_ < 0.001). CC32 also reflected a univariate pattern of SA (92%), with minor contributions from curvature (5%) and CT (2%), indicating that children with lower birth weight and/or being born premature show bi-directional patterns of smaller and larger occipital and parietal SA.

Low Birth Weight and Prematurity showed a positive association with CC37 (t = 5.71, Cohen’s D = 0.14, p_adjusted_ < 0.001). CC37 reflected a multivariate pattern of CT (63%), SA (28%) and curvature (9%) indicating that children with lower birth weight and/or being born premature have reduced SA and increased CT in the insular cortex, and increased SA and reduced CT in the frontal cortex.

Low Birth Weight and Prematurity showed a negative association with CC39 (t = −5.23, Cohen’s D = −0.13, p_adjusted_ < 0.001). CC39 showed a univariate pattern of SA (91%), with some contributions from curvature (6%) and CT (3%) indicating that children with a low birth weight and/or being born premature have patterns of bi-directional differences in frontal SA.

### Newborn Birth Complications

Linear mixed effects modelling revealed significant associations between Newborn Birth Complications and CC10 and CC16 (Tab. 2 and Fig. 3).

Newborn Birth Complications showed a positive association with CC10 (t = 3.52, Cohen’s D = 0.08, p_adjusted_ = 0.023), which reflected a univariate pattern of SA (91%), with smaller contributions from curvature (6%) and CT (3%), indicating that children who experienced complications at birth have smaller occipital SA and larger temporal, parietal, and frontal SA at about 10 years of age.

Newborn Birth Complications also showed a negative association with CC16 (t = −4.41, Cohen’s D = −0.10, p_adjusted_ < 0.001), indicating that beyond the previously described results, children who experienced birth complications have larger temporal and insular SA, smaller frontal SA, and a pattern of reduced CT spread across cortical regions.

## Discussion

The developing fetus is highly sensitive to its intrauterine environment and events occurring around the time of birth. In this study, we employed a multifactorial and multivariate fusion imaging approach to investigate how a diverse range of pregnancy- and birth-related risk factors are associated with variation in cortical morphometry in late childhood. Our findings showed that children whose mothers experienced somatic health complications during pregnancy, as well as children with low birth weight and premature birth, exhibited smaller global SA. Additionally, children experiencing low birth weight, premature birth and birth complications demonstrated distinct regional cortical patterns.

Our findings linking maternal pregnancy complications to smaller global SA of the child align with previous ABCD Study findings investigating either a single or a few pregnancy-related variables and regional cortical measures (3, 6, 9), as well as a previous multifactor multivariate fusion study (28). Smaller human imaging studies and animal studies have also linked maternal pregnancy complications to altered brain structure in the offspring (9, 44–47). Our results also support prior research linking low birth weight and prematurity with reduced global or widespread SA in children (5, 28, 35–40, 42). A longitudinal study comparing term-born and preterm-born children showed that the difference in SA is mostly established at birth, and that most cortical regions displayed similar SA developmental trajectories over time (39). Further, our multivariate fusion approach of cortical data allowed us to move beyond well-established associations between pregnancy- and birth-related risk factors and unique SA effects, revealing additional regional patterns across different morphometrics.

Our findings align with previous studies that reported no associations between low birth weight/prematurity or newborn birth complications and global CT (5, 37, 42). Instead, we observed regional CT variations in children experiencing low birth weight and/or prematurity (36– 38). Specifically, our multivariate patterns support previous studies linking low birth weight and prematurity to increased frontal (CC7, CC16) (38, 39), reduced temporal (CC7, CC16, CC37) (36, 39), reduced pre-and post-central (CC16) (37), and increased occipital (CC37) (37, 39) CT. However, other studies report more widespread, different, or contradictory patterns of regional CT (36–39). In contrast, our findings differ from studies reporting no regional associations with newborn birth complications (28, 42, 48). Reasons for these discrepancies could be methodological differences such as different reduction methods (28), and lower sample size (42, 48).

Overall, our results indicate that maternal pregnancy complications, low birth weight and prematurity have global homogenous effects on child SA, while the same factors in addition to birth complications have regional heterogenous effects on child CT (39). Premature birth will cause the fetus to miss the later part of pregnancy, a critical period for in-utero SA expansion (13, 36). In contrast, the peak growth of CT occurs mid-gestation, which could explain some of the divergent effects observed on CT compared to SA.

Our analysis revealed no significant associations between maternal substance use and later child cortical structure, which contrasts previous studies (7, 8, 10, 29–34). This discrepancy may be due to several factors, including relatively low variance in several of the maternal substance use variables in the ABCD Study sample, reliance on self-reported data that may have led to underreporting, and that we were unable to account for dose, frequency, and timing of substance use. However, potential associations may have gone undetected, as we did not explore indirect mediation effects. Previous research suggests that maternal substance use can influence fetal size (49, 50), which may subsequently impact the child’s cortex –an association that we observed. Finally, although speculative, multifactorial data-driven reduction methods could mask potential specific effects.

Contrary to previous studies on cortical folding (6, 28, 36), we found minimal relations between pregnancy- and birth-related risk factors on child cortical curvature. This discrepancy is likely partly due to our cortical components having limited contributions from curvature, thereby restricting our ability to identify such associations from the outset. Moreover, differences in the measures of cortical folding and other methodological aspects could also account for the discrepant findings.

While our cross-sectional design is not sensitive to changes in cortical morphology over time, our findings align with known developmental patterns. SA peaks around age 11 before declining (13), and since our sample (ages 9–11) likely has not reached this peak, the observed smaller SA may reflect either slower development or a consistently smaller SA across childhood. A recent study reported no significant differences in SA trajectories between term- and very preterm-born children from birth to age 13 (39), suggesting that SA differences are established at birth and remain stable. In contrast, CT follows more variable developmental patterns. Consistent with our findings, prior research shows that CT differences between preterm- and term-born children can widen, shrink, or remain stable over time (39).

Although caution is warranted when interpreting neurobiological maturity based on cross-sectional findings, cortical differences in late childhood associated with pregnancy- and birth-related risk factors have been suggested to reflect brain maturity (3–6, 9, 37, 39). For instance, smaller global SA has been associated with lower general cognitive ability in children born with very low birth weight (37), reduced CT in inferior frontal and medial parietal regions has been associated with impaired language and memory function in pre-term born children (39), and CT in middle frontal gyrus has been shown to mediate the association between prematurity and children’s language function (51). These previous findings could imply that our results showing associations between low birth weight and prematurity and smaller global SA and regional patterns of altered CT may also reflect functional outcomes, although this remains speculative.

There are several potential mechanisms that can explain the observed associations between pregnancy- and birth-related risk factors and child cortical structure. One such mechanism is fetal exposure to maternal stress during pregnancy (52, 53). Proxies for maternal stress such as prenatal depression have been associated with pregnancy complications (54, 55), preterm birth (56, 57), and child cortical structure (58, 59). Fetal exposure to maternal stress may impair the barrier of the placenta by affecting an enzyme (11beta-hydroxysteroid dehydrogenase type 2 (11β-HSD2)) involved in stress regulation, allowing for more stress hormones to reach the fetus (60, 61). Indeed, reduced activity of 11β-HSD2 has been associated with incidences of both pregnancy- and birth-related risk factors (62–64).

Another potential mechanism is hypoxia-related neural harm (42, 65), changes in number of neurons (37, 66), and accelerated neuronal maturation (67). Hypoxia is the lack of sufficient oxygen to the baby’s brain during delivery (68), that could be indicated by the variables “oxygen treatment”, “blue at birth” and “no breath at birth” in the present study. Regional differences in cortical associations with hypoxia may be explained by the location of different cerebral vessels (42, 65).

Other underlying mechanisms could be shared genetic susceptibility between pregnancy- and birth-related risk factors and child cortical morphometry (36, 69, 70) and epigenetic effects. For instance, DNA methylation has been shown to meditate the association between birth weight and cortical development (35). Moreover, maternal circadian disruptions during pregnancy have been associated with variations in placental methylation, indicating an alteration of the intrauterine environment (71), which could affect fetal neurodevelopment. Lastly, maternal inflammation during pregnancy has been associated with both birth-related risk factors (72, 73) and neurodevelopment in the child (74, 75).

The current study has several limitations. First, data on pregnancy- and birth-related risk factors were obtained from retrospective self-reporting, which can introduce both recall biases and response biases. Second, our observational study does not allow us to determine whether the pregnancy- and birth-related risk factors causally drive the observed cortical effects, or if other complex relationships are at play. Third, while considering a wide array of pregnancy- and birth-related risk factors offers a comprehensive perspective, it comes at the expense of not addressing specific details such as the timing and dose of substance use and severity and timing of pregnancy- and birth-related complications (42, 76–78). Finally, the ABCD Study sample is not ethnically representative of the US population (or the world), with nearly 80% of participants identifying as white (79). Similarly, participants have higher socioeconomic status than the general US population (80). Finally, given the minimal amount of missing data, the challenge of keeping the dimension-reduction pipeline consistent across multiple imputed data sets, and the absence of standard pooling methods for permutation-based p-values, we opted for single stochastic imputation. This approach does not capture between-imputation variance, and the model’s standard errors may therefore be slightly underestimated. A detailed discussion on how limitations might have impacted our findings is presented in SI Appendix, Discussion, Limitations and Future Research.

Our study revealed that maternal somatic health during pregnancy as well as low birth weight and/or prematurity in children were associated with unique global SA signatures later in the child’s life. Low birth weight, prematurity and experiencing birth complications were also associated with shared regional morphological patterns. While specific neurobiological mechanisms remain to be explored, our findings highlight the importance of supporting mothers and their unborn babies during early-life critical periods to foster optimal child development.

## Materials and Methods

### Sample

The data was drawn from the ABCD Study, a dataset comprising nearly 12.000 children aged 9-11 years, along with their parents. This ongoing longitudinal study tracks brain development and child health across the US, with data collected from 21 research centers located across the country (https://abcdstudy.org/). The current study used data from the baseline assessment, release 4.0. Of the 11.876 participants, 9758 had pre-processed, quality controlled vertex-wise imaging data (81). From these, 1327 children were excluded due to their ABCD Developmental History Questionnaire (43) being completed by their non-biological mother. An additional 35 participants were excluded due to having more than 15% missing responses for the pregnancy- and birth-related variables of interest. This yielded a final sample of 8396 participants aged 8.9 – 11.1 years (mean = 9.9, SD = 0.6), including 582 twins (291 pairs) and 2558 siblings. Sample demographics are presented in SI Appendix section “Methods”, SI Appendix Fig. S8 and SI Appendix Tab. S2.

### Assessment of Pregnancy- and Birth-Related Variables

Pregnancy- and birth-related variables were collected through retrospective reporting from the biological mother using the ABCD Developmental History Questionnaire (43). From this questionnaire, 23 pregnancy variables and 12 birth variables were identified. An overview of included variables is provided in SI Appendix Tab. S3, with further details about questions provided in SI Appendix Table S4 and recoding of variables in SI Appendix section “Methods”. A correlation matrix for the variables is shown in SI Appendix Fig. S9. As expected, several of the variables showed near-zero variance as identified using the “nearZeroVar” function from the caret-package (version 6.0-92) in R, version 4.2.1 (https://www.r-project.org/). Despite potentially offering less information about the dependent variable and concerns of overfitting, we decided to keep all selected variables as rare extreme factors (e.g. maternal use of oxycontin and maternal infection of rubella during pregnancy) could be highly influential on cortical structure, and as they could be aggregated into overarching factors, thereby amplifying overall variance. Results from analyses excluding variables with near-zero variance (SI Appendix Tab. S5) yielded similar results (SI Appendix Results, Fig. S10-S12 and Tab. S6-S7). Missing pregnancy- and birth-related values were imputed using the “mice” package (version 3.14.0) in R. We applied predictive mean matching for numeric variables and logistic regression imputation for binary variables. Imputation was performed in five iterations, and we selected the first for analysis. Continuous variables, birth weight and weeks born before due date, were z-standardized.

We used the “MCA” and “FAMD” functions from the FactoMineR package (version 2.4) in R to reduce and extract underlying factors from the pregnancy- and birth-related variables, which were split accordingly. MCA was applied to the binary pregnancy variables, while FAMD handled the mixed binary and continuous birth variables. This reduction yielded several dimensions, and we chose two for each set of variables for further analyses, which were z-standardized. We assessed imputation validity by comparing our resulting four dimensions to the same dimensions generated using only individuals with complete datapoints, yielding highly converging results (SI Appendix Fig. S13-S14).

### MRI Acquisition, Preprocessing, and Quality Control

The MRI data was acquired on 29 different 3T scanners from Siemens Prisma, Philips, and General Electric (GE) 750 (82). T1-weighted images were obtained through an inversion-prepared RF-spoiled gradient echo scan, featuring 1-millimeter isotropic voxel resolution, and was consistently performed as the second sequence. An adult-sized coil was used. Further details regarding the MRI acquisition, including parameters for the scans, are provided elsewhere (82). The T1-weighted images underwent an initial quality control by the ABCD-team, checking for quality issues such as incorrect acquisition parameters, imaging artifacts, or corrupted data files, using both automated and manual methods (83). 367 participants were excluded due to not passing this binary include/exclude quality control (81, 83).

We processed the T1-weighted images using FreeSurfer 7.1 (https://surfer.nmr.mgh.harvard.edu/) (84, 85). The present study used vertex-wise calculations of cortical morphology (84), specifically CT, SA, and curvature (see SI Appendix section “Methods” for details). 104 participants were excluded due to missing surface data (81). FreeSurfer also calculates the quality metric Euler number, which reflects the total amount of holes in the initial surface (86). 64 participants were excluded due to having surface holes =< 200 (81). CT, SA, and curvature surface maps were mapped to FreeSurfer’s standard space *fsaverage* and smoothed with a Gaussian kernel of 15mm full width at half maximum (FWHM) to increase signal and contrast. Finally, neuroComBat in R was used to harmonize the data across the 29 scanners to adjust for systematic and unwanted scanner related variance (87). Age, sex, twin and sibling status, the pregnancy-related dimensions (Maternal Pregnancy Complications and Maternal Substance Use), and the birth-related dimensions (Low Birth Weight and Prematurity and Newborn Birth Complications) were included as covariates in the harmonization procedure. Box plots of cortical measures pre‐and post‐neuroCombat are presented in SI Appendix Fig. S15-S17.

### Multivariate Fusion of Cortical Metrics

CT, SA, and curvature were fused using FMRIB’s Linked-ICA (41). FLICA can reduce and fuse large amounts of data into independent components across different metrics, even if the input has different smoothness and signal- and contrast to noise. Linked-ICA finds shared patterns across metrics but can also capture univariate patterns when present (41, 88). In the current study, we employed Linked-ICA with 1000 iterations and explored model orders ranging from 20 to 50 components. Logarithmic transformation was applied exclusively to the SA data.

We chose a model order of 40 cortical components based on the cophenetic correlation coefficient (SI Appendix Fig. S18) and visual inspection of the surface maps, attempting to reach a pragmatic balance between data reduction, interpretability and bi-hemispheric specificity. A cophenetic correlation coefficient represents how well the original data structure is being captured at the chosen number of components (81). One component was driven by a one single subject (SI Appendix Fig. S19) and was excluded from further analyses, yielding a final number of 39 cortical components.

### Statistical Analysis

All statistical analyses were conducted in R. Associations between the pregnancy- and birth-related dimensions and each of the cortical components were tested by linear mixed effects modelling using the “lmer” function from the lme4-package (version 1.1-29). The “lme-dscore” function from the EMAtools-package (version 0.1.4) was used to calculate Cohen’s D. The 39 cortical components were dependent variables, while the four pregnancy- and birth-related dimensions were independent variables in separate models, resulting in a total of 156 models. In all models, age and sex were added as fixed effects, while family status (coded with a unique value for each family) and zygosity (with each monozygotic twin-pair receiving an identical value) was modelled as random effects to address dependency in the data. All continuous variables not yet z-standardized were then z-standardized. Due to the non-normality of residuals in parametric analysis (SI Appendix Tab. S8), significance testing was performed using permutation of the independent variable (5000 iterations). P-values were calculated as the proportion of permuted t-values exceeding the observed t-value and were adjusted for multiple comparisons using the Hochberg procedure, applied separately for each pregnancy- and birth-related dimension (FEW <0.05). R code is available online (https://osf.io/xeufj/).

### Ethical Considerations

Each ABCD site is responsible for following ethical guidelines related to implementation of the study, and each site has developed site-specific standard procedures for the implementation of these guidelines (89, 90). All procedures were approved by a central Institutional Review Board (IRB) at the University of California, San Diego (89), while an External Advisory Board is monitoring ethical issues (91). Parental and child informed consent have been obtained for all participants (89, 90). The present study was conducted in line with the Declaration of Helsinki and was approved by the Regional Committee for Medical and Health Research Ethics (REK 2019/943).

## Supporting information

SI_appendix_Linking_Pregnancy_and_Birth_Related_Factors_to_a_Multivariate_Fusion_of_Child_Cortical_Structure

## Acknowledgments

Data used in the preparation of this article were obtained from the ABCD Study (https://abcdstudy.org), held in the NIMH Data Archive. This is a multisite, longitudinal study designed to recruit more than 10,000 children aged 9-10 and follow them over 10 years into early adulthood. The ABCD Study® is supported by the National Institutes of Health and additional federal partners under award numbers U01DA041048, U01DA050989, U01DA051016, U01DA041022, U01DA051018, U01DA051037, U01DA050987, U01DA041174, U01DA041106, U01DA041117, U01DA041028, U01DA041134, U01DA050988, U01DA051039, U01DA041156, U01DA041025, U01DA041120, U01DA051038, U01DA041148, U01DA041093, U01DA041089, U24DA041123, U24DA041147. A full list of supporters is available at https://abcdstudy.org/federal-partners.html. A listing of participating sites and a complete listing of the study investigators can be found at https://abcdstudy.org/consortium_members/. ABCD consortium investigators designed and implemented the study and/or provided data but did not necessarily participate in the analysis or writing of this report. This manuscript reflects the views of the authors and may not reflect the opinions or views of the NIH or ABCD consortium investigators. This work was funded by the Research Council of Norway (#288083, #300767, #323951), the South-Eastern Norway Regional Health Authority (#2021070, #2023012, #500189), and the European Research Council under the European Union’s Horizon 2020 research and Innovation program (ERC StG Grant No. 802998).

