## Supplementary material for "Linking Pregnancy- and Birth-Related Risk Factors to a Multivariate Fusion of Child Cortical Structure": SI_appendix_Linking_Pregnancy_and_Birth_Related_Factors_to_a_Multivariate_Fusion_of_Child_Cortical_Structure

**This PDF file includes:**

Supporting text
Figures S1 to S19
Tables S1 to S8
SI References

### Supporting Information Text

#### Methods

**Calculations of Parental Education and Family Income.** Parental education was assessed by "What is the highest grade or level of school you have completed or the highest degree you have received?" and "What is the highest grade or level of school your partner completed or highest degree they received?". The highest score was chosen. The scale was 0 = Never attended/Kindergarten only; 1 = 1st grade; 2 = 2nd grade; 3 = 3rd grade; 4 = 4th grade; 5 = 5th grade; 6 = 6th grade; 7 = 7th grade; 8 = 8th grade; 9 = 9th grade; 10 = 10th grade; 11 = 11th grade; 12 = 12th grade; 13 = High school graduate; 14 = GED or equivalent; 15 = Some college; 16 = Associate degree: Occupational; 17 = Associate degree: Academic Program; 18 = Bachelor's degree; 19 = Master's degree; 20 = Professional School degree (ex. MD); 21 = Doctoral degree (ex. PhD). Family income was assessed by the question "What is your total combined family income for the past 12 months?". The scale was 1= Less than $5,000; 2=$5,000 through $11,999; 3=$12,000 through $15,999; 4=$16,000 through $24,999; 5=$25,000 through $34,999; 6=$35,000 through $49,999; 7=$50,000 through $74,999; 8= $75,000 through $99,999; 9=$100,000 through $199,999; 10=$200,000 and greater. See distributions of parental education and family income in SI Appendix Fig. S8.

**Calculations of Ethnicity**. Ethnicity was assessed by parental report through "What race do you consider the child to be? Please check all that apply." Asian consists of Asian Indian, Chinese, Filipino, Japanese, Korean, Vietnamese, and Other Asian. AIAN is American Indian Native or Alaska Native. NHOPI is Native Hawaiian, Guamanian, Samoan, and Other Pacific Islander. See SI Appendix Tab S2.

**Recoding of Pregnancy-and Birth-Related Variables.** The following recoding of pregnancy- and birth-related variables was performed. To indicate substance use during pregnancy, the use of "prescriptive medications", "tobacco", "alcohol", "marijuana", "cocaine / crack", "heroine / morphine", "oxycontin" and "other drugs" was coded as "yes" irrespective of the individuals awareness of their pregnancy status. Caffeine use during pregnancy was coded as "yes" if use was reported "at least once a day", "less than once a day but more than once a week", "less than once a week," or if "caffeine" was marked under "other drugs". The variable birth weight was converted from pounds (lbs) to kilograms. Within the variable "weeks born before due date" "0" indicated that the child was born at its due date in week 40, while "1" that the child was born one week early i.e. within week 39, and so on. As the specific week of birth was not specified for children born 12 or more weeks premature, these values were recoded to 15.5, representing the midpoint between 27 and 22 weeks. The cutoff of 22 was chosen as it currently signifies the earliest week at which a child is likely to survive a premature birth (1).

**Vertex-Wise Calculations of Cortical Morphology**. Cortical thickness was calculated as the shortest vertex-wise distance between the white (inner) and the pial (outer) surface, while vertex-wise surface area was calculated by summing the areas of the triangles that converge at each specific vertex on the white surface (2, 3). Curvature was measured as the average of the two primary curvature directions at each vertex (2). In this context, a higher value implies a decrease in the radius of curvature, often further indicating more pronounced folds, or sulci and gyri.

**Reduction of Pregnancy- and Birth-Related Variables**. The selection of dimensions for further analyses was guided by visual inspection of an "elbow" plot of explained variance. The point at which the curve began to flatten, combined with a pragmatic decision to capture as much variance as possible while reducing the data to a limited number of overarching dimensions, informed the final choice (SI Appendix Fig. S1).

Of note, in the dimension space for Low Birth Weight and Prematurity, birth weight was indicated having a positive relationship with the dimension score (SI Appendix Fig. S3) which is counter intuitive, and not in line with previous studies (4–6). Nevertheless, reduction techniques like MCA and FAMD capture complex relationships amongst the variables and reflect the effect of the variables combined. Therefore, interpreting a single variable within a dimension is not necessarily ideal. Indeed, separate correlation analyses of Low Birth Weight and Prematurity and birth weight yielded a negative association (SI Appendix Fig. S9), indicating that in the present sample, individuals with a lower birth weight loaded higher on the Low Birth Weight and Prematurity dimension.

### Results

**Near-Zero Variance.** Results from analyses excluding variables with near-zero variance at a rate of >0.99 for the most common to the second most common response are shown in SI Appendix Fig. S10-S12 and SI Appendix Tab. S6-S7. The findings showed significant associations between Maternal Pregnancy Complications and CC1, Maternal Substance Use and CC31 and CC34, Low Birth Weight and Prematurity and CC1, CC7, CC16, CC18, CC32, CC37 and CC39, and Newborn Birth Complications and CC10, CC16 and CC18.

### Discussion

**Limitations and Future Research.** The current study has several limitations. First, data on pregnancy- and birth-related risk factors were obtained from retrospective self-reporting, which can introduce both recall biases (i.e. forgetting, temporal inaccuracies, and memory alterations due to emotional events related to pregnancy and birth) and response biases (i.e. modifying responses based on social norms). For instance, low variance in the variable related to prenatal substance use could partly reflect both forgetting and intentional underreporting, thereby limiting the signal available for tests on child cortical structure. This, in turn, may explain our null finding for this factor. Future studies should aim to mitigate this by using objective biomarkers in real time, such as urine samples during pregnancy. Moreover, a high-risk sample would increase overall users. Second, our observational design does not allow us to determine whether the pregnancy- and birth-related risk factors causally drive the observed cortical effects. Third, while considering a wide array of pregnancy- and birth-related risk factors offers a comprehensive perspective, it comes at the expense of addressing specific details such as timing and dose of substance use, or severity and timing of pregnancy- and birth-related complications (7–10). For instance, our null findings for substance use could be different if we specifically tested high dosage of drugs with early exposure in pregnancy. An intensive, prospective study design could probe timing- and dosage-specific aspects. Finally, the ABCD sample is not ethnically representative of the US population (or the world), with nearly 80% of participants identifying as white (11). Ethnic health disparities in the US, particularly pertaining to African American women are well documented (12–14). Also, participants have higher socioeconomic status than the general US population (15). It is possible that the associations between pregnancy- and birth-related risk factors and child brain structure could yield additional findings with a larger non-white and lower SES sample, as this likely would include more individuals with higher risk.

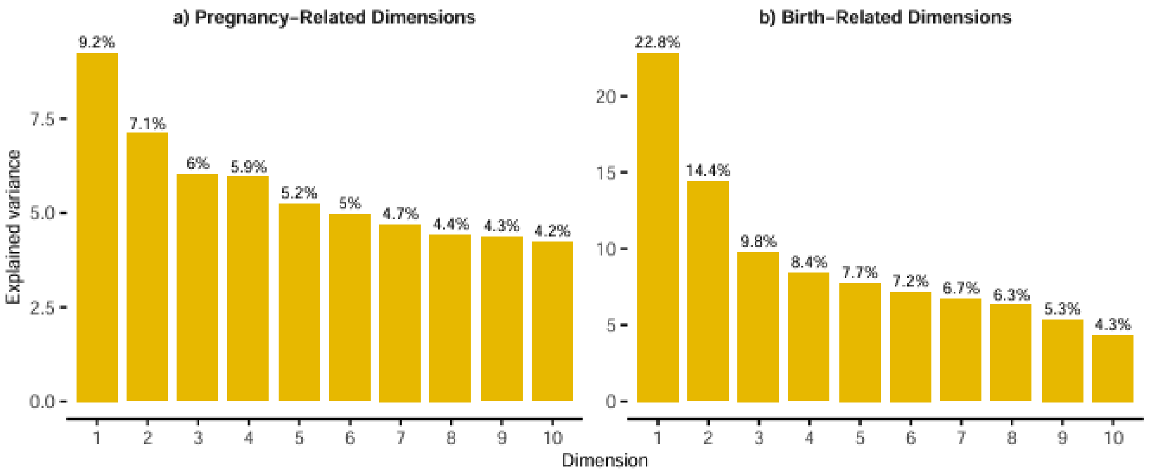


**Fig. S1.** Dimensions from the reduction analysis. The figure shows explained percentage variance of the first 10 dimensions.

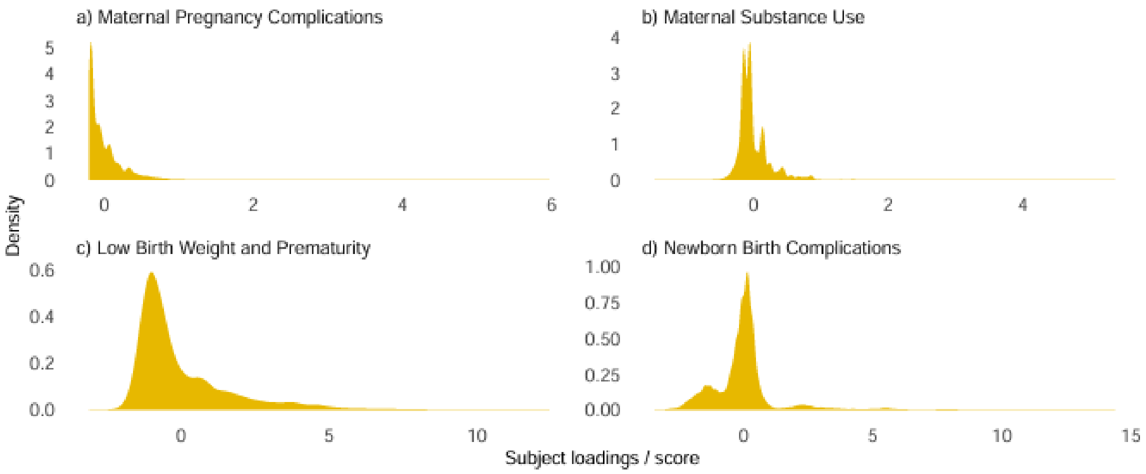


**Fig. S2.** Density plots for pregnancy- and birth-related dimensions. The figure shows density plots of the distribution of subject loadings/scores for the dimensions.

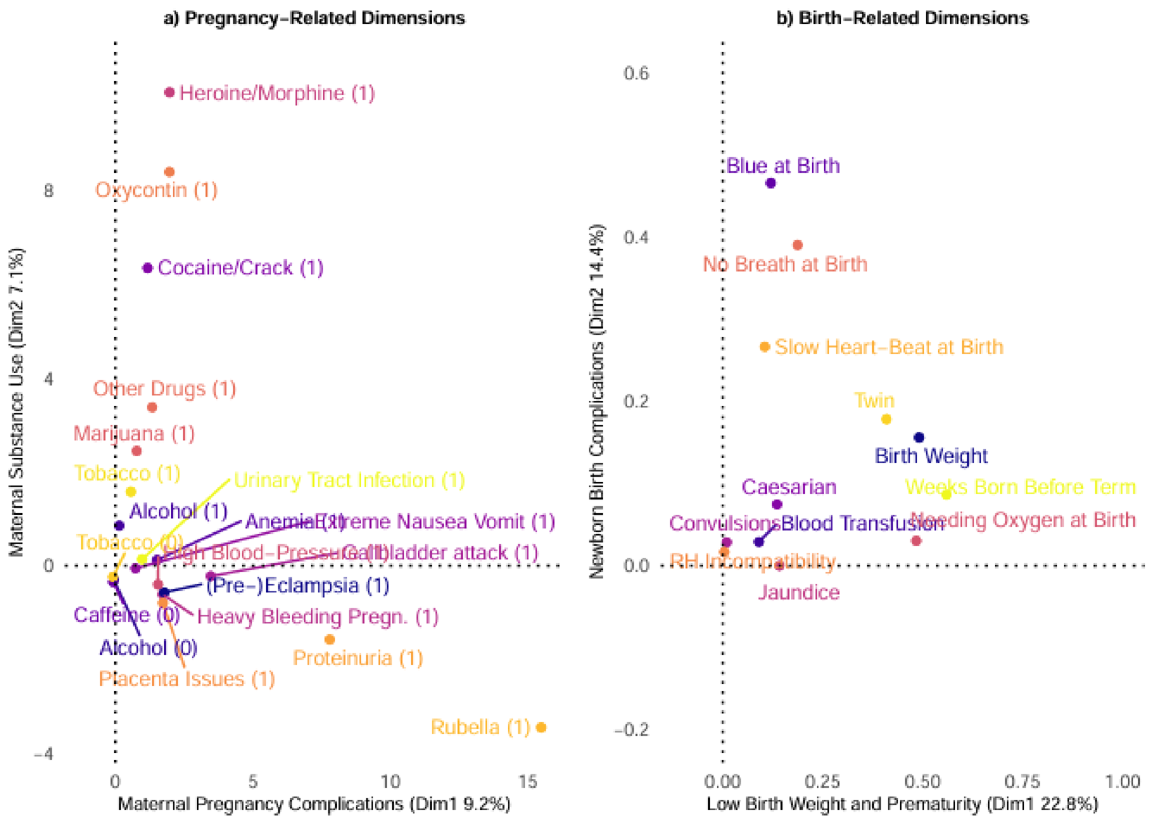


**Fig. S3.** Contributing variables and the dimensional spaces from the reduction analysis**.** The figure shows the variables with the highest contributions and their coordinates in the dimensional space for the a) pregnancy variables and the b) birth variables. Percentages represent amount of total explained variance in the for pregnancy- and birth-related data separately. (1) and (0) is a response of yes and no, respectively. Figure a) shows the 10 top contributing variables for the dimensions Maternal Pregnancy Complications and Maternal Substance Use.

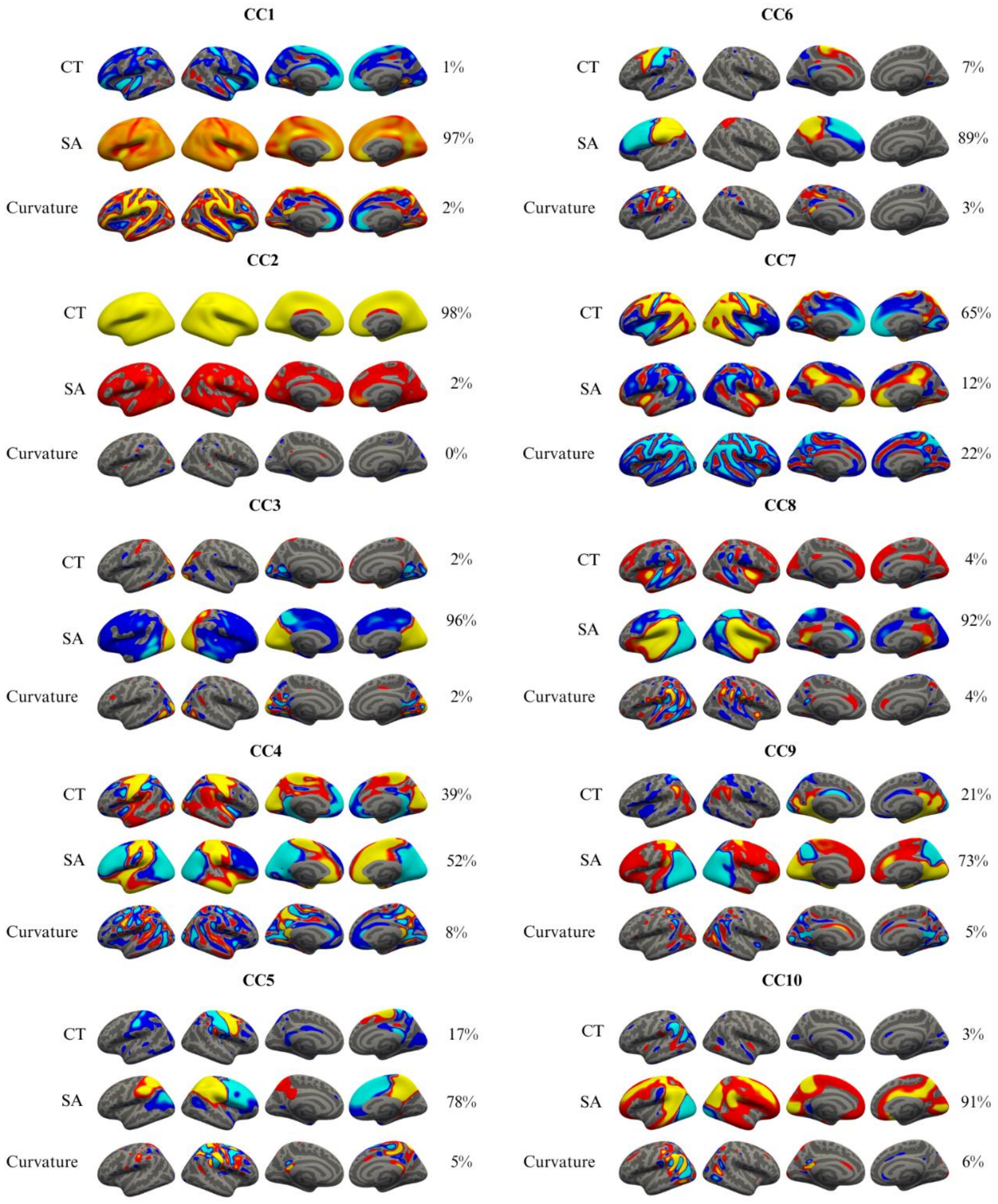


**Fig. S4.** Surface maps for the cortical components (CC1 – CC10)**.**

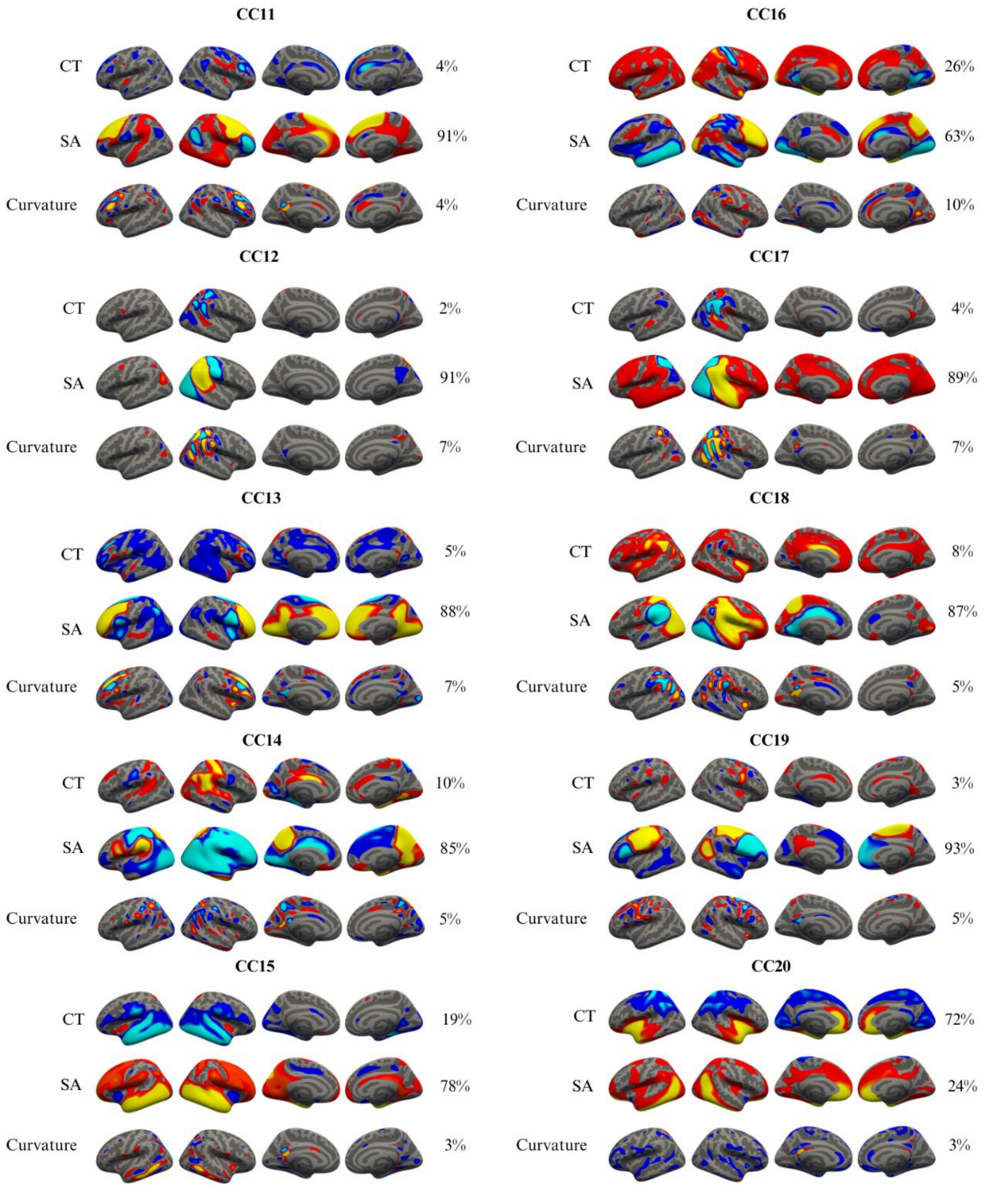


**Fig. S5.** Surface maps for the cortical components (CC11 – CC20)**.**

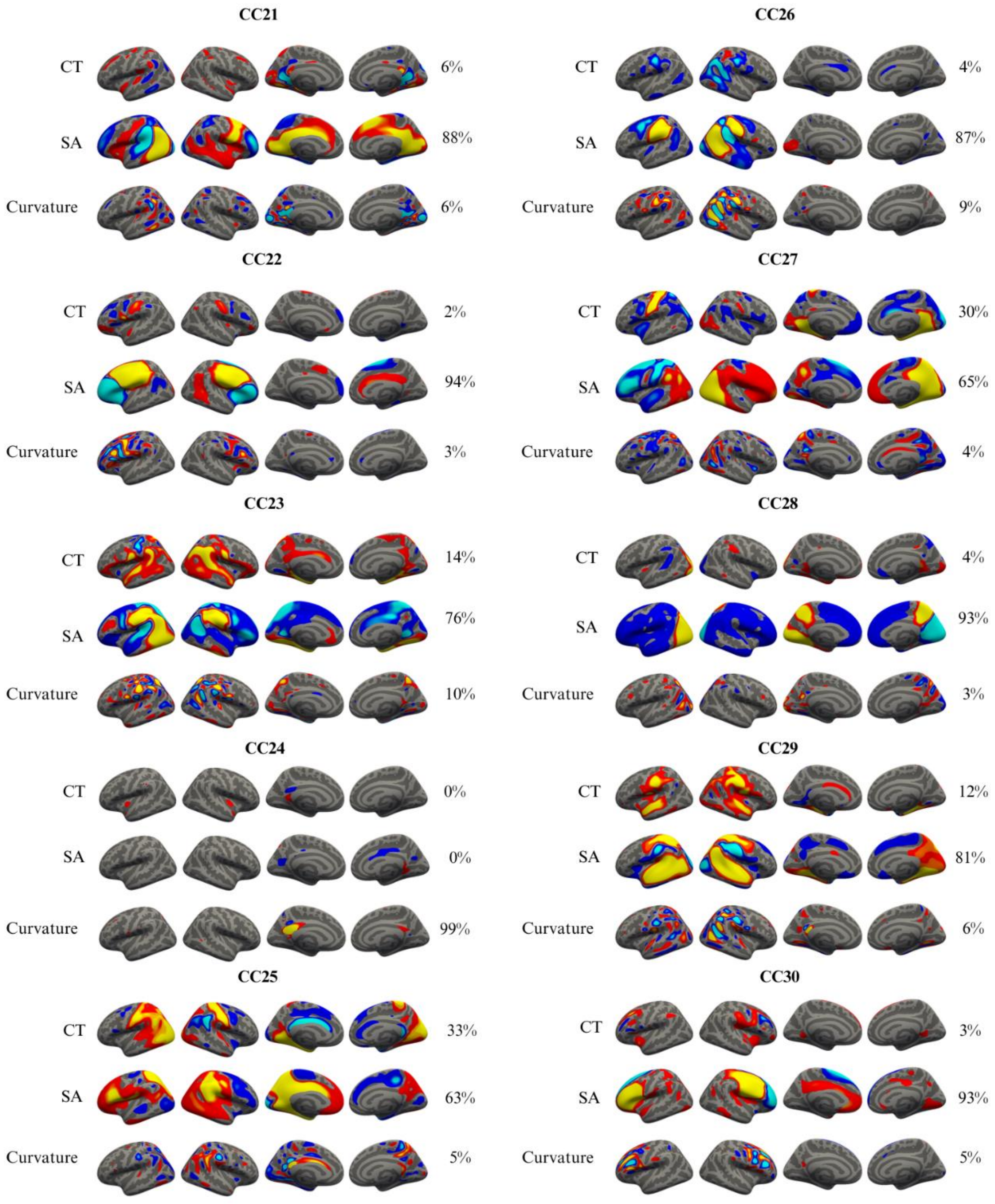


**Fig. S6.** Surface maps for the cortical components (CC21 – CC30). CC24 was excluded from the statistical analysis due to being driven by mainly one single subject.

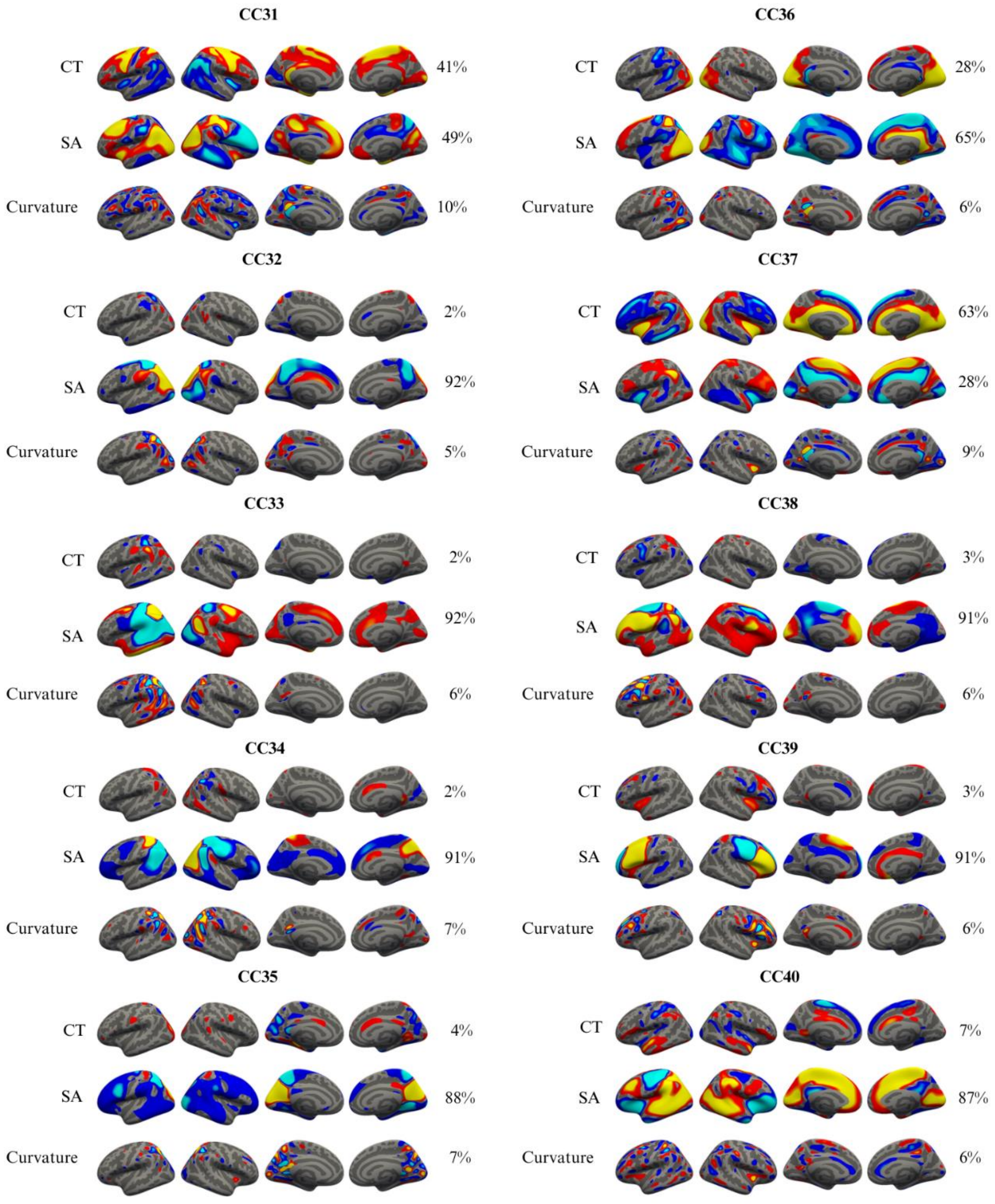


**Fig. S7**. Surface maps for the cortical components (CC31 – CC40).

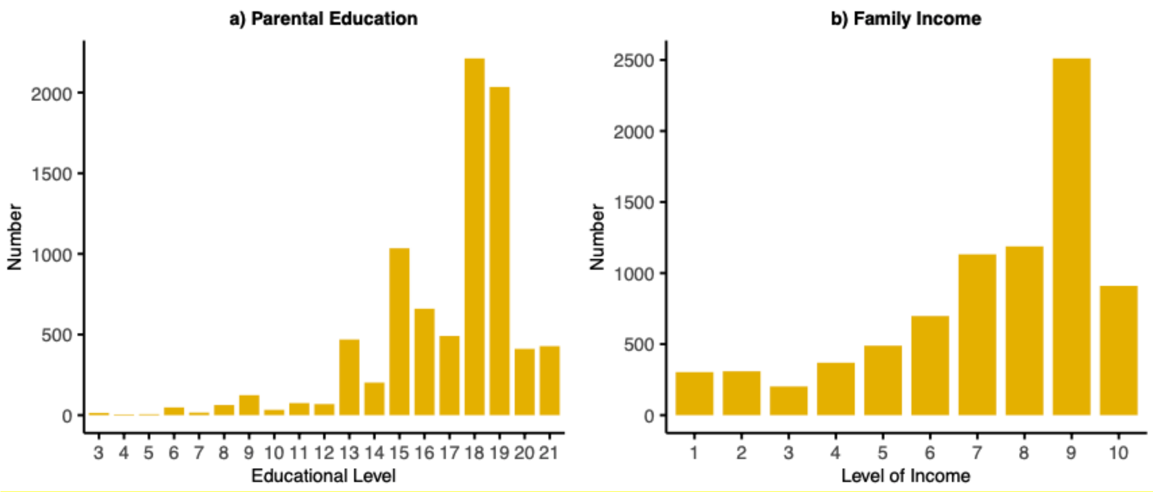


**Fig. S8.** Parental education and family income. The figure shows the distribution of parental education and family income within the final sample for those with such data available.

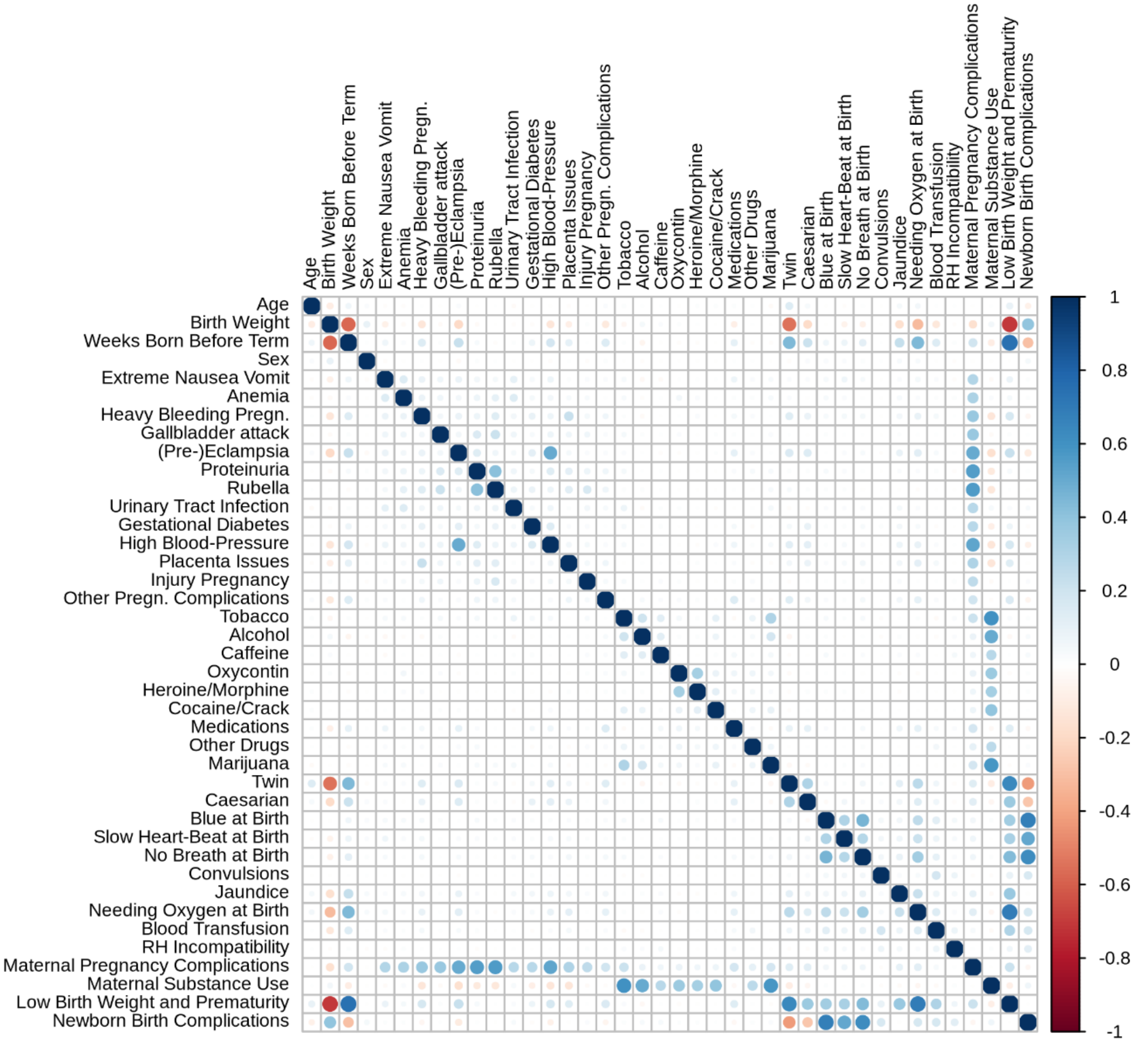


**Fig. S9.** Correlation matrix. The figure shows a Pearson's correlation matrix for age, sex, pregnancy- and birth-related variables, and pregnancy- and birth-related dimensions from the reduction analyses.

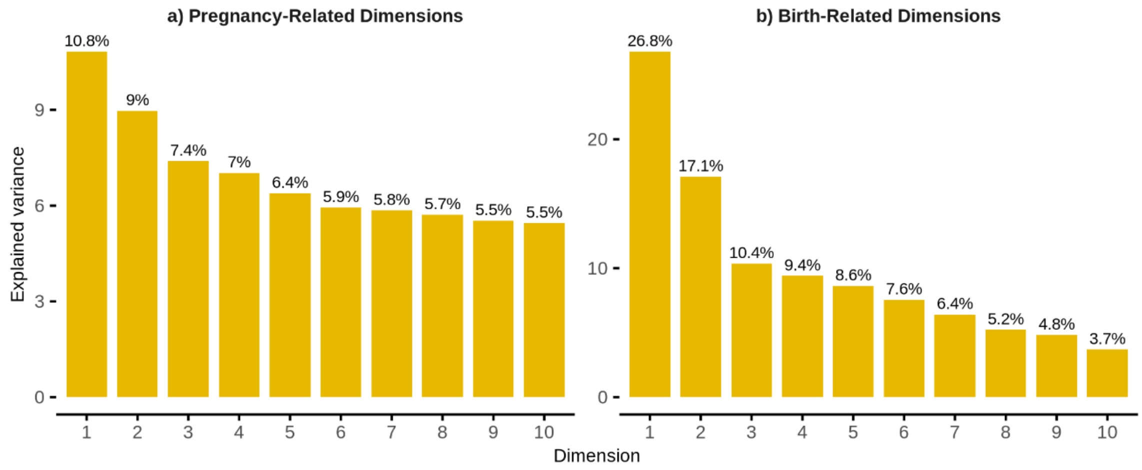


**Fig. S10.** Dimensions from the reduction analysis using an imputed dataset excluding variables with near-zero variance. The figure shows explained percentage variance of the first 10 dimensions.

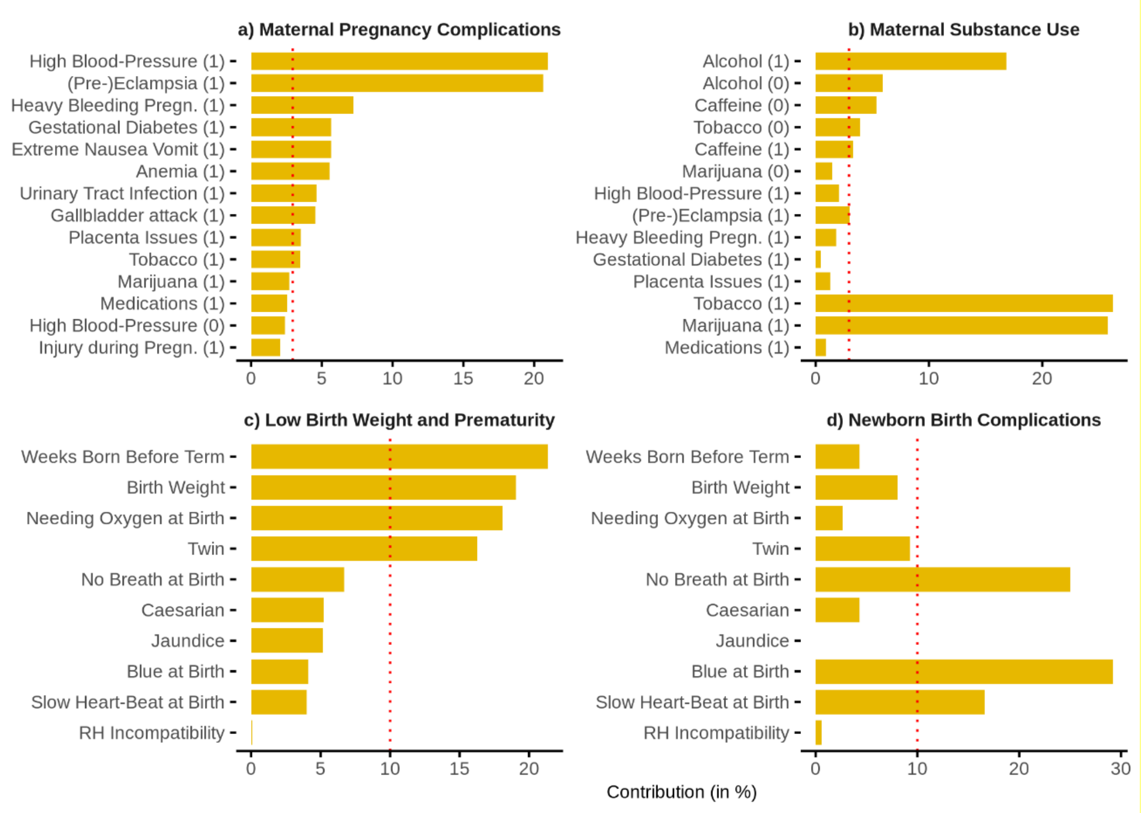


**Fig. S11.** Variable contribution to pregnancy- and birth-related dimensions using an imputed dataset excluding variables with near-zero variance. The figure shows the variables with the highest contributions to each dimension. The x-axis indicates the percentage contribution while the red dotted line represents the expected average contribution if the variables were uniform. a) and b) are pregnancy dimensions while c) and d) are birth dimensions. (1) and (0) indicates whether the response to the variable is yes or no, respectively. Figure a) and b) show the top 14 contributing variables in the given dimension.

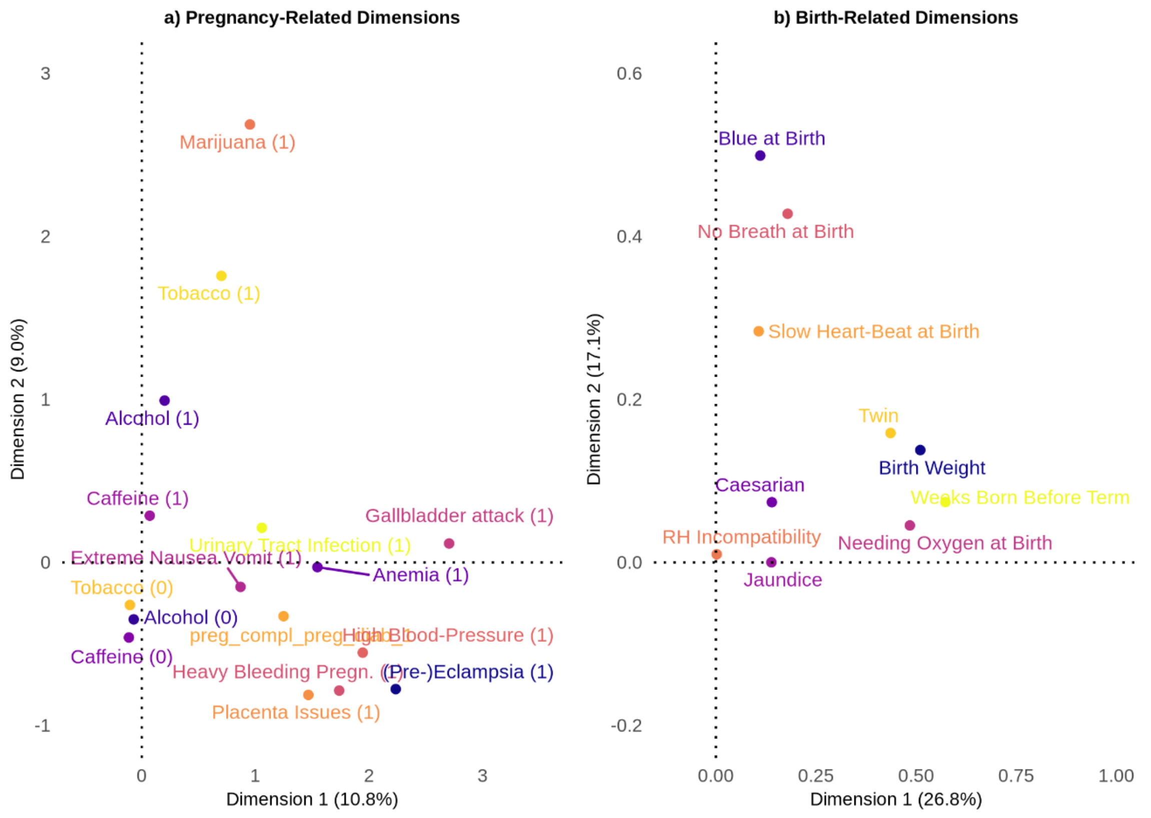


**Fig. S12.** Contributing variables and the dimensional spaces from the reduction analysis, using an imputed dataset excluding variables with near-zero variance**.** The figure shows the variables with the highest contributions and their coordinates in the dimensional space for the a) pregnancy variables and the b) birth variables. Percentages represent amount of total explained variance in the for pregnancy- and birth-related data separately. (1) and (0) is a response of yes and no, respectively. Figure a) shows the 10 top contributing variables for the dimensions Maternal Pregnancy Complications and Maternal Substance Use.

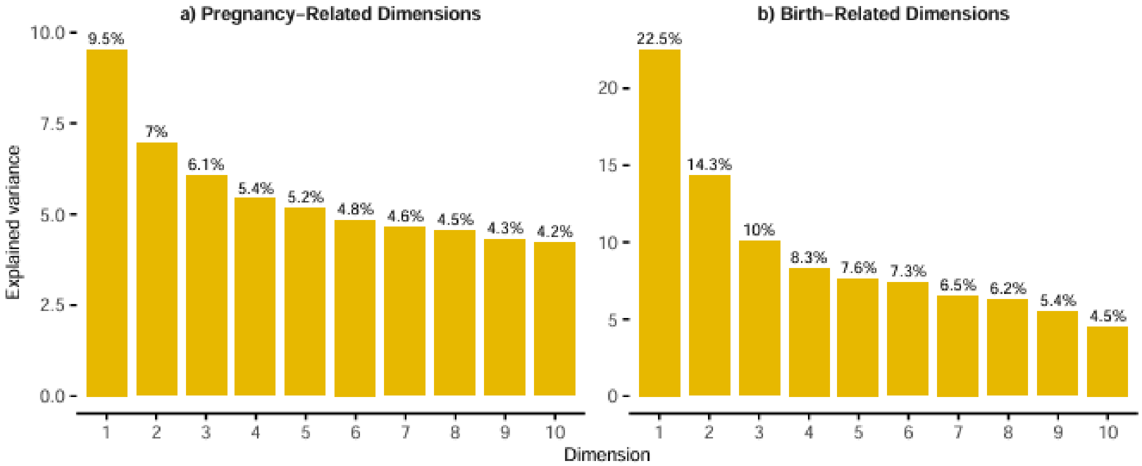


**Fig. S13.** Dimensions from the reduction analysis using a dataset with dropped NAs instead of using imputation. The figure shows explained percentage variance of the first 10 dimensions.

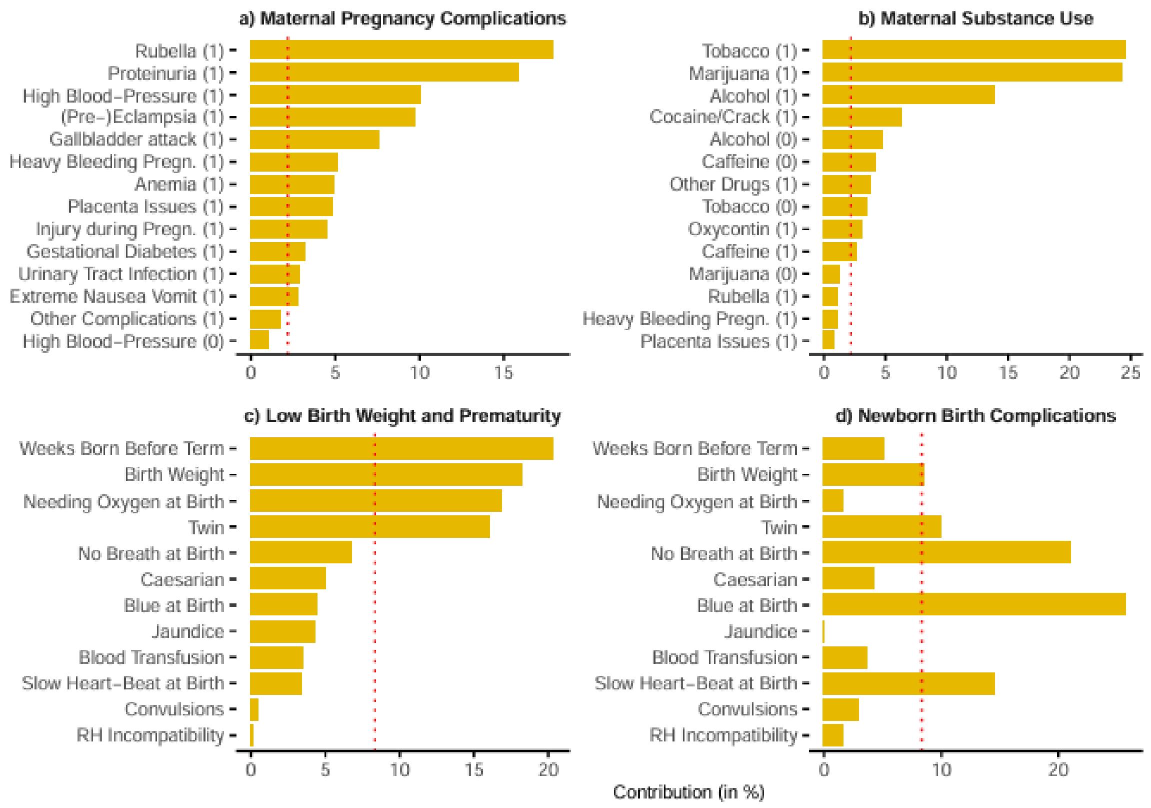


**Fig. S14.** Variable contribution to pregnancy- and birth-related dimensions using a dataset with dropped NAs instead of using imputation. The figure shows the variables with the highest contributions to each dimension. The x-axis indicates the percentage contribution while the red dotted line represents the expected average contribution if the variables were uniform. a) and b) are pregnancy dimensions while c) and d) are birth dimensions. (1) and (0) indicates whether the response to the variable is yes or no, respectively. Figure a) and b) show the top 14 contributing variables in the given dimension.

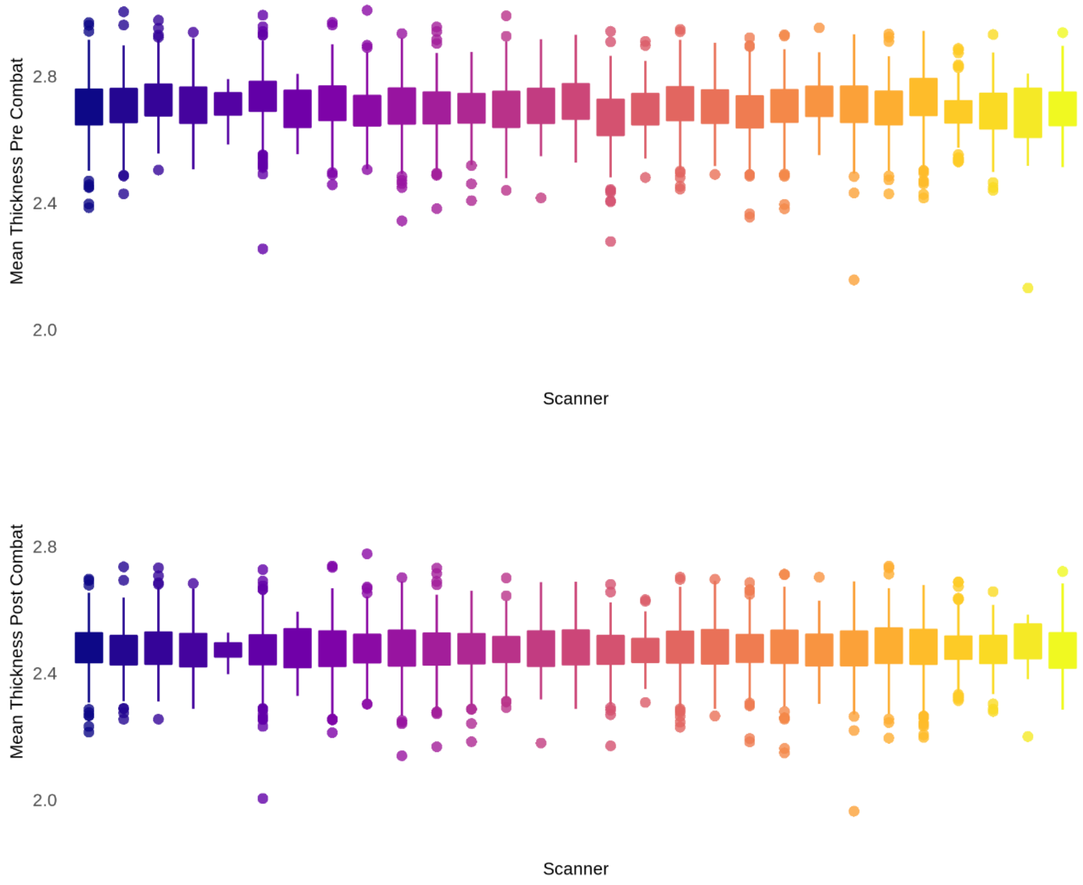


**Fig. S15.** Boxplots showing mean cortical thickness pre and post scanner harmonization using neuroCombat.

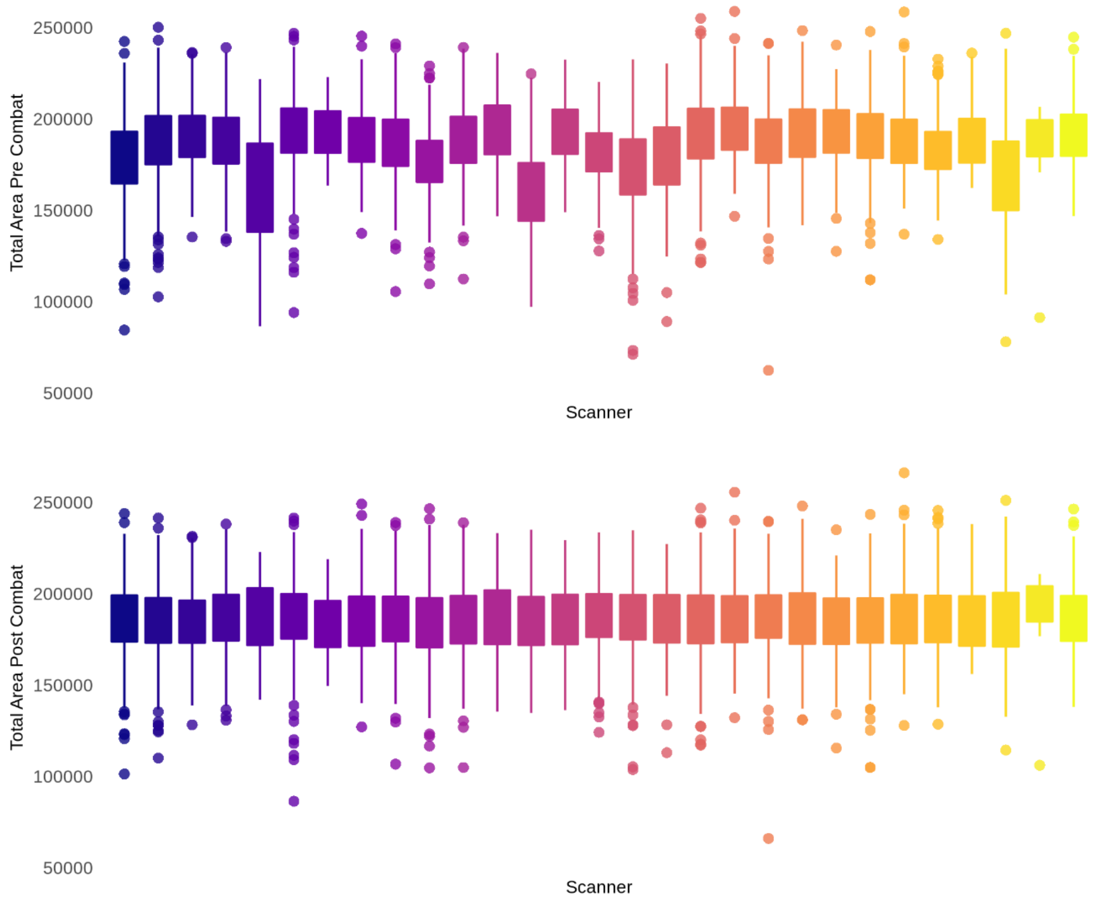


**Fig. S16.** Boxplots showing total surface area pre and post scanner harmonization using neuroCombat.

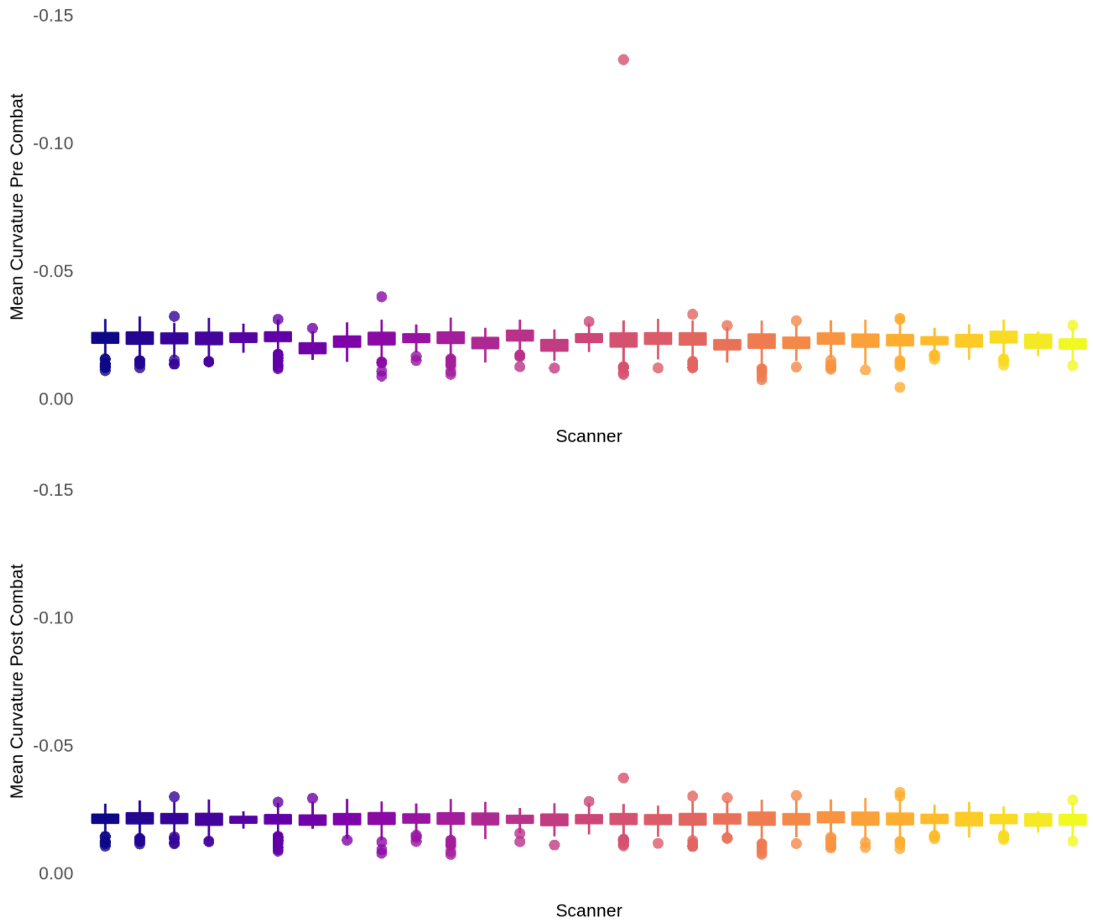


**Fig. S17.** Boxplots showing mean curvature pre and post scanner harmonization using neuroCombat.

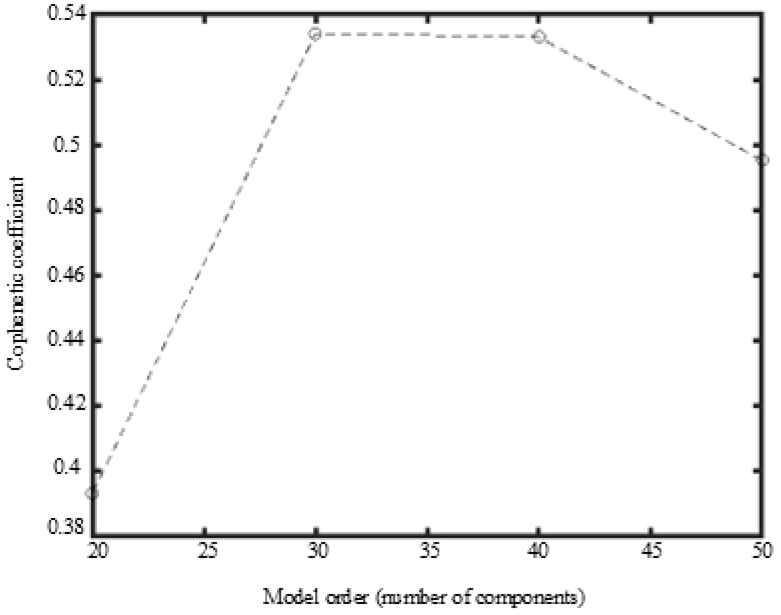


**Fig. S18.** Linked Independent Component Analysis Model Order Selection. The figure shows the cophenetic correlation coefficient for all the fusion model orders tested; with 20, 30, 40, and 50 cortical components.

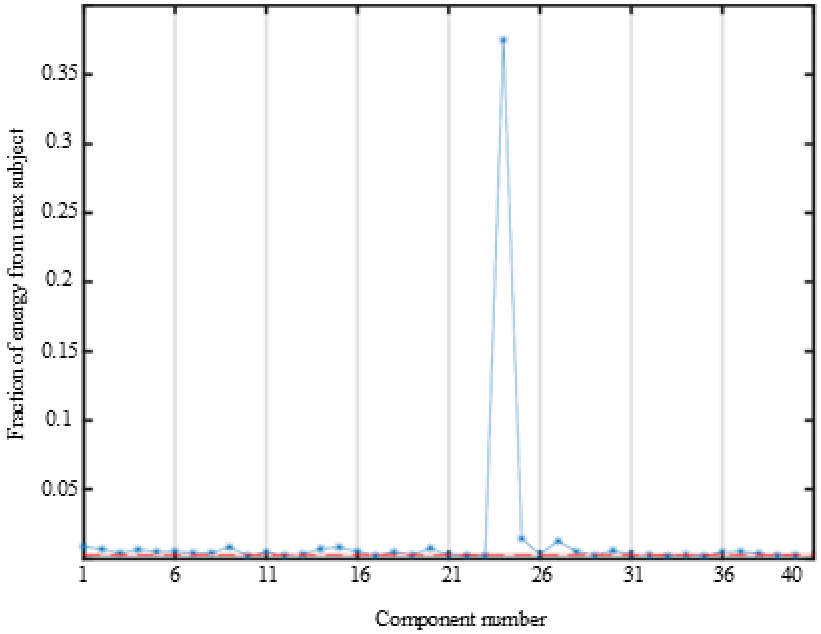


**Fig. S19.** The figure shows the proportion of the energy (subject weight squared) that comes from one participant. Higher values will indicate that the component is driven by one or a few outlier subjects.

**Table S1.** Weighting of cortical morphology and explained variance for the cortical components.

| Cortical Component | Total Explained variance | Weighting | | |
|---|---|---|---|---|
| | | CT | SA | Curvature |
| 1 | 27.80 % | 1 % | 97 % | 2 % |
| 2 | 6.86 % | 98 % | 2 % | 0 % |
| 3 | 5.11 % | 2 % | 96 % | 2 % |
| 4 | 4.02 % | 39 % | 52 % | 8 % |
| 5 | 3.75 % | 17 % | 78 % | 5 % |
| 6 | 2.56 % | 7 % | 89 % | 3 % |
| 7 | 2.32 % | 65 % | 12 % | 22 % |
| 8 | 2.30 % | 4 % | 92 % | 4 % |
| 9 | 2.15 % | 21 % | 73 % | 5 % |
| 10 | 2.07 % | 3 % | 91 % | 6 % |
| 11 | 1.94 % | 4 % | 91 % | 4 % |
| 12 | 1.80 % | 2 % | 91 % | 7 % |
| 13 | 1.71 % | 5 % | 88 % | 7 % |
| 14 | 1.69 % | 10 % | 85 % | 5 % |
| 15 | 1.57 % | 19 % | 78 % | 3 % |
| 16 | 1.53 % | 26 % | 63 % | 10 % |
| 17 | 1.51 % | 4 % | 89 % | 7 % |
| 18 | 1.50 % | 8 % | 87 % | 5 % |
| 19 | 1.50 % | 3 % | 93 % | 5 % |
| 20 | 1.49 % | 72 % | 24 % | 3 % |
| 21 | 1.45 % | 6 % | 88 % | 6 % |
| 22 | 1.45 % | 2 % | 94 % | 3 % |
| 23 | 1.44 % | 14 % | 76 % | 10 % |
| 24 | 1.43 % | 0 % | 0 % | 99 % |
| 25 | 1.39 % | 33 % | 63 % | 5 % |
| 26 | 1.36 % | 4 % | 87 % | 9 % |
| 27 | 1.34 % | 30 % | 65 % | 4 % |
| 28 | 1.31 % | 4 % | 93 % | 3 % |
| 29 | 1.27 % | 12 % | 81 % | 6 % |
| 30 | 1.25 % | 3 % | 93 % | 5 % |
| 31 | 1.24 % | 41 % | 49 % | 10 % |
| 32 | 1.22 % | 2 % | 92 % | 5 % |

| | | | | |
|---|---|---|---|---|
| 33 | 1.17 % | 2 % | 92 % | 6 % |
| 34 | 1.13 % | 2 % | 91 % | 7 % |
| 35 | 1.12 % | 4 % | 88 % | 7 % |
| 36 | 1.11 % | 28 % | 65 % | 6 % |
| 37 | 1.11 % | 63 % | 28 % | 9 % |
| 38 | 1.06 % | 3 % | 91 % | 6 % |
| 39 | 1.00 % | 3 % | 91 % | 6 % |
| 40 | 0.98 % | 7 % | 87 % | 6 % |

The table shows the weighting for each cortical measure (cortical thickness, surface area, and curvature) and the explained variance for each cortical component in the cortical decomposition.

**Table S2.** Sample demographics.

| | | Number (%) | Mean (SD) | Range |
|---|---|---|---|---|
| Total N | | 8396 | | |
| Age, years | | | 9.92 (0.63) | |
| Age, years | | | | 8.9 - 11.1 |
| Sex | | | | |
| | Female | 4018 (47.9) | | |
| | Male | 4378 (52.1) | | |
| Siblings | | 2558 (30.5) | | |
| Twins | | 582 (6.9) | | |
| Ethnicity | | | | |
| | White | 6525 (77.7) | | |
| | Black / African American | 1686 (20.1) | | |
| | Other | 508 (6.1) | | |
| | Asian | 415 (4.9) | | |
| | AIAN | 260 (3.1) | | |
| | NHOPI | 47 (0.6) | | |

The table shows sample demographics for the sample. SD = standard deviation. N = sample size. AIAN = American Indian Native or Alaska Native. NHOPI = Native Hawaiian, Guamanian, Samoan, and Other Pacific Islander.

**Table S3.** Descriptive summary of pregnancy- and birth-related variables.

| Pregnancy- and birth-related variables | Number (%) | | Mean (SD) | Range |
|---|---|---|---|---|
| | Yes | No | | |
| Use prescriptive medications | 1565 (18.6) | 6831 (81.4) | | |
| Use prenatal vitamins | 8069 (96.1) | 327 (3.9) | | |
| Use caffeine | 5145 (61.3) | 3251 (38.7) | | |
| Use tobacco | 1088 (13.0) | 7308 (87.0) | | |
| Use alcohol | 2177 (25.9) | 6219 (74.1) | | |
| Use marijuana | 450 (5.4) | 7946 (94.6) | | |
| Use cocaine / crack | 32 (0.4) | 8364 (99.6) | | |
| Use heroin / morphine | 9 (0.1) | 8387 (99.9) | | |
| Use oxycontin | 16 (0.2) | 8380 (99.8) | | |
| Use other drugs | 51 (0.6) | 8345 (99.4) | | |
| Severe nausea and vomiting | 1154 (13.7) | 7242 (86.3) | | |
| Heavy bleeding | 371 (4.4) | 8025 (95.6) | | |
| Pre-eclampsia, eclampsia, or toxemia | 637 (7.6) | 7759 (92.4) | | |
| Gall bladder attack | 96 (1.1) | 8300 (98.9) | | |
| Persistent proteinuria | 42 (0.5) | 8354 (99.5) | | |
| Rubella during 1st trimester | 11 (0.1) | 8385 (99.9) | | |
| Severe anemia | 356 (4.2) | 8040 (95.8) | | |
| Urinary tract infections | 640 (7.6) | 7756 (92.4) | | |
| Pregancy-related diabetes | 563 (6.7) | 7833 (93.3) | | |
| Pregnancy-related high blood pressure | 859 (10.2) | 7537 (89.8) | | |
| Previa, abruptio, or other problems with placenta | 251 (3.0) | 8145 (97.0) | | |
| Accident or injury requiring medical care | 155 (1.8) | 8241 (98.2) | | |
| Other condition requiring medical care | 765 (9.1) | 7631 (90.9) | | |
| Twin birth | 1696 (20.2) | 6700 (79.8) | | |
| Caesarian | 3208 (38.2) | 5188 (61.8) | | |
| Blue at birth | 282 (3.4) | 8114 (96.6) | | |
| Slow heart beat | 251 (3.0) | 8145 (97.0) | | |
| Did not breathe at first | 410 (4.9) | 7986 (95.1) | | |
| Convulsions | 11 (0.1) | 8385 (99.9) | | |
| Jaundice needing treatment | 1440 (17.2) | 6956 (82.8) | | |
| Required oxygen | 869 (10.4) | 7527 (89.6) | | |
| Required blood transfusion | 42 (0.5) | 8354 (99.5) | | |
| Rh incompatibility | 244 (2.9) | 8152 (97.1) | | |
| Birth weight in kg. | | | 2.99 (0.66) | |
| Birth weight in kg. | | | | 0.9 - 6.35 |
| Weeks born before due date | | | 0.96 (2.27) | |

| | |
|---|---|
| Weeks born before due date | 0 - 15.5 |

SD = standard deviation.

**Table S4.** Questions on pregnancy- and birth-related variables from the questionnaire

| Element name | Question | Answers | Variable in present study |
|---|---|---|---|
| devhx_8_prescript_med | Before the biological mother/you found out she was pregnant, but while she/you might have been pregnant with this child, did she/you use any of the following? Prescription medications? | 1 = Yes; 0 = No; 999 = Don't know | Use prescriptive medications |
| devhx_9_prescript_med | Once you / biomom knew you/she were pregnant, were you/biomom using any of the following? Prescription medications? | 1 = Yes; 0 = No; 999 = Don't know | Use prescriptive medications |
| devhx_10 | Did you/biological mother take prenatal vitamins during the pregnancy? | 1 = Yes; 0 = No; 999 = Don't know; -1 = Not applicable | Use prenatal vitamins |
| devhx_caffeine_11 | Did you/biological mother have any caffeine during pregnancy (from conception until delivery)? | 1 = Yes - at least once a day; 2 = Yes - less than once a day but more than once a week; 3 = Yes - less than once a week; 0 = No; 999 = Don't know; -1 = Not applicable | Use caffeine |
| devhx_8_tobacco | Before knowing of pregnancy. Tobacco? | 1 = Yes; 0 = No; 999 = Don't know | Use tobacco |
| devhx_9_tobacco | Knowing of pregnancy. Tobacco? | 1 = Yes; 0 = No; 999 = Don't know | Use tobacco |
| devhx_8_alcohol | Before knowing of pregnancy. Alcohol? | 1 = Yes; 0 = No; 999 = Don't know | Use alcohol |
| devhx_9_alcohol | Knowing of pregnancy. Alcohol? | 1 = Yes; 0 = No; 999 = Don't know | Use alcohol |
| devhx_8_marijuana | Before knowing of pregnancy. Marijuana? | 1 = Yes; 0 = No; 999 = Don't know | Use marijuana |
| devhx_9_marijuana | Knowing of pregnancy. Marijuana? | 1 = Yes; 0 = No; 999 = Don't know | Use marijuana |
| devhx_8_coc_crack | Before knowing of pregnancy. Cocaine/Crack? | 1 = Yes; 0 = No; 999 = Don't know | Use cocaine / crack |
| devhx_9_coc_crack | Knowing of pregnancy. Cocaine/Crack? | 1 = Yes; 0 = No; 999 = Don't know | Use cocaine / crack |
| devhx_8_her_morph | Before knowing of pregnancy. Heroin/Morphine? | 1 = Yes; 0 = No; 999 = Don't know | Use heroine / morphine |
| devhx_9_her_morph | Knowing of pregnancy. Heroin/Morphine? | 1 = Yes; 0 = No; 999 = Don't know | Use heroine / morphine |
| devhx_8_oxycont | Before knowing of pregnancy. Oxycontin? | 1 = Yes; 0 = No; 999 = Don't know | Use oxycontin |
| devhx_9_oxycont | Knowing of pregnancy. Oxycontin? | 1 = Yes; 0 = No; 999 = Don't know | Use oxycontin |

| | | | |
|---|---|---|---|
| devhx_8_other_drugs | Before knowing of pregnancy. Any other drugs? | 1 = Yes; 0 = No; 999 = Don't know | Use other drugs |
| devhx_8_other1 _name_2 | Before knowing of pregnancy. Drug 1 | 0 = None; 1 = Amphetamines or methamphetamine (meth; 13 = Barbituates; 2 = Benzodiazepines; 3 = Caffeine; 4 = Cathinones (bath salts; 5 = Fake or synthetic marijuana (like spice or K2); 6 = GHB (liquid G or Georgia home boy); 7 = Hallucinogens (LSD or acid; 8 = Inhalants; 9 = Ketamine (special K); 10 = MDMA (ecstasy; 11 = Opioids; 12 = Other; 999 = Don't Know | Use other drugs |
| devhx_9_other_drugs | Knowing of pregnancy. Any other drugs? | 1 = Yes; 0 = No; 999 = Don't know | Use other drugs |
| devhx_9_other1 _name_2 | Knowing of pregnancy. Drug 1 | 0 = None; 1 = Amphetamines or methamphetamine (meth; 13 = Barbituates; 2 = Benzodiazepines; 3 = Caffeine; 4 = Cathinones (bath salts; 5 = Fake or synthetic marijuana (like spice or K2); 6 = GHB (liquid G or Georgia home boy); 7 = Hallucinogens (LSD or acid; 8 = Inhalants; 9 = Ketamine (special K); 10 = MDMA (ecstasy; 11 = Opioids; 12 = Other; 999 = Don't Know | Use other drugs |
| devhx_10a3_p | During the pregnancy with this child, did you/biological mother have any of the following conditions? Severe nausea and vomiting extending past the 6th month or accompanied by weight loss? | 1 = Yes; 0 = No; 999 = Don't know | Severe nausea and vomiting |

| | | | |
|---|---|---|---|
| devhx_10b3_p | During the pregnancy with this child, did you/biological mother have any of the following conditions? Heavy bleeding requiring bed rest or special treatment? | 1 = Yes; 0 = No; 999 = Don't know | Heavy bleeding |
| devhx_10c3_p | During the pregnancy with this child, did you/biological mother have any of the following conditions? Pre-eclampsia, eclampsia, or toxemia? | 1 = Yes; 0 = No; 999 = Don't know | Pre-eclampsia, eclampsia, or toxemia |
| devhx_10d3_p | During the pregnancy with this child, did you/biological mother have any of the following conditions? Severe gall bladder attack? | 1 = Yes; 0 = No; 999 = Don't know | Gall bladder attack |
| devhx_10e3_p | During the pregnancy with this child, did you/biological mother have any of the following conditions? Persistent proteinuria? | 1 = Yes; 0 = No; 999 = Don't know | Persistent proteinuria |
| devhx_10f3_p | During the pregnancy with this child, did you/biological mother have any of the following conditions? Rubella (German measles) during first 3 months of pregnancy? | 1 = Yes; 0 = No; 999 = Don't know | Rubella during 1st trimester |
| devhx_10g3_p | During the pregnancy with this child, did you/biological mother have any of the following conditions? Severe anemia? | 1 = Yes; 0 = No; 999 = Don't know | Severe anemia |
| devhx_10h3_p | During the pregnancy with this child, did you/biological mother have any of the following conditions? Urinary tract infections? | 1 = Yes; 0 = No; 999 = Don't know | Urinary tract infections |
| devhx_10i3_p | During the pregnancy with this child, did you/biological mother have any of the following conditions? Pregnancy-related diabetes? | 1 = Yes; 0 = No; 999 = Don't know | Pregancy-related diabetes |
| devhx_10j3_p | During the pregnancy with this child, did you/biological mother have any of the following conditions? Pregnancy-related high blood pressure? | 1 = Yes; 0 = No; 999 = Don't know | Pregnancy-related high blood pressure |
| devhx_10k3_p | During the pregnancy with this child, did you/biological mother have any of the following conditions? Previa, abruptio, or other problems with the placenta? | 1 = Yes; 0 = No; 999 = Don't know | Previa, abruptio, or other problems with placenta |

| | | | |
|---|---|---|---|
| devhx_10l3_p | During the pregnancy with this child, did you/biological mother have any of the following conditions? An accident or injury requiring medical care? | 1 = Yes; 0 = No; 999 = Don't know | Accident or injury requiring medical care |
| devhx_10m3_p | During the pregnancy with this child, did you/biological mother have any of the following conditions? Any other conditions requiring medical care? | 1 = Yes; 0 = No; 999 = Don't know | Other condition requiring medical care |
| birth_weight_lbs | Birth weight pounds | | Birth weight in kg (converted) |
| devhx_12a_p | Was the child born prematurely? | 1 = Yes; 0 = No; 999 = Don't know | Weeks born before due date |
| devhx_12_p | About how many weeks premature was the child when they were born? | 1 = 1; 2 = 2; 3 = 3; 4 = 4; 5 = 5; 6 = 6; 7 = 7; 8 = 8; 9 = 9; 10 = 10; 11 = 11; 12 = 12; 13 = Greater than 12; 999 = Don't know | Weeks born before due date |
| devhx_5_p | Does your child have a twin? | 1 = Yes; 0 = No; 999 = Don't know | Twin birth |
| devhx_13_3_p | Was your child born by Caesarian section? | 1 = Yes; 0 = No; 999 = Don't know | Caesarian |
| devhx_14a3_p | Did he/she have any of the following complications at birth? Blue at birth? | 1 = Yes; 0 = No; 999 = Don't know | Blue at birth |
| devhx_14b3_p | Did he/she have any of the following complications at birth? Slow heart beat? | 1 = Yes; 0 = No; 999 = Don't know | Slow heart beat |
| devhx_14c3_p | Did he/she have any of the following complications at birth? Did not breathe at first? | 1 = Yes; 0 = No; 999 = Don't know | Did not breathe at first |
| devhx_14d3_p | Did he/she have any of the following complications at birth? Convulsions? | 1 = Yes; 0 = No; 999 = Don't know | Convulsions |
| devhx_14e3_p | Did he/she have any of the following complications at birth? Jaundice needing treatment? | 1 = Yes; 0 = No; 999 = Don't know | Jaundice needing treatment |
| devhx_14f3_p | Did he/she have any of the following complications at birth? Required oxygen? | 1 = Yes; 0 = No; 999 = Don't know | Required oxygen |
| devhx_14g3_p | Did he/she have any of the following complications at birth? Required blood transfusion? | 1 = Yes; 0 = No; 999 = Don't know | Required blood transfusion |
| devhx_14h3_p | Did he/she have any of the following complications at birth? Rh incompatibility? | 1 = Yes; 0 = No; 999 = Don't know | Rh incompatibility |

The table shows the questions used in the retrospective reporting from the mothers. For a full overview of variables, see the ABCD Developmental History Questionnaire (16).

**Table S5.** Pregnancy- and birth-related variables with zero or near-zero variance.

| Frequency cut | |
|---|---|
| >0.95 | >0.99 |
| Use prenatal vitamins | Persistent proteinuria |
| Heavy bleeding | Rubella during 1st trimester |
| Gall bladder attack | Convulsions |
| Persistent proteinuria | Required blood transfusion |
| Rubella during 1st trimester | Use cocaine / crack |
| Severe anemia | Use heroin / morphine |
| Previa, abruptio, or other problems with placenta | Use oxycontin |
| Accident or injury requiring medical care in pregnancy | Use other drugs |
| Blue at birth | |
| Slow heart beat | |
| Did not breathe at birth | |
| Convulsions | |
| Required blood transfusion | |
| Rh incompatibility | |
| Use cocaine / crack | |
| Use heroin / morphine | |
| Use oxycontin | |
| Use other drugs | |

The table shows pregnancy- and birth-related variables with a ratio of either >0.95 or >0.99 for the most common to the second most common response.

**Table S6.** Associations between pregnancy-related dimensions and cortical components from sensitivity analysis excluding variables with near-zero variance.

| Cortical Component | Maternal Pregnancy Complications | | | | Maternal Substance Use | | | |
|---|---|---|---|---|---|---|---|---|
| | Cohen´s D | t-value | p-value (unadj.) | p-value (adj.) | Cohen´s D | t-value | p-value (unadj.) | p-value (adj.) |
| 1 | -0.11 | -4.75 | **<0.001** | **<0.001** | 0.05 | 2.04 | **0.042** | 0.972 |
| 2 | -0.07 | -3.01 | **0.003** | 0.101 | 0.01 | 0.30 | 0.766 | 0.972 |
| 3 | -0.01 | -0.46 | 0.647 | 0.958 | 0.05 | 2.00 | **0.045** | 0.972 |
| 4 | -0.02 | -0.79 | 0.432 | 0.958 | -0.03 | -1.32 | 0.185 | 0.972 |
| 5 | -0.01 | -0.59 | 0.557 | 0.958 | 0.02 | 0.89 | 0.371 | 0.972 |
| 6 | 0.02 | 0.64 | 0.525 | 0.958 | 0.01 | 0.44 | 0.663 | 0.972 |
| 7 | -0.04 | -1.47 | 0.142 | 0.958 | 0.00 | 0.07 | 0.942 | 0.972 |
| 8 | 0.01 | 0.55 | 0.582 | 0.958 | -0.01 | -0.48 | 0.631 | 0.972 |
| 9 | 0.00 | -0.06 | 0.954 | 0.958 | -0.05 | -2.18 | **0.030** | 0.972 |
| 10 | -0.06 | -2.57 | **0.010** | 0.371 | 0.02 | 0.90 | 0.367 | 0.972 |
| 11 | -0.02 | -0.89 | 0.372 | 0.958 | 0.02 | 0.67 | 0.501 | 0.972 |
| 12 | 0.04 | 1.56 | 0.119 | 0.958 | 0.00 | -0.04 | 0.972 | 0.972 |
| 13 | -0.02 | -0.78 | 0.434 | 0.958 | -0.04 | -1.72 | 0.085 | 0.972 |
| 14 | -0.03 | -1.31 | 0.191 | 0.958 | 0.02 | 0.89 | 0.375 | 0.972 |
| 15 | -0.06 | -2. 39 | **0.017** | 0.569 | -0.03 | -1.19 | 0.234 | 0.972 |
| 16 | 0.03 | 1.39 | 0.166 | 0.958 | -0.04 | -1.63 | 0.103 | 0.972 |
| 17 | 0.01 | 0.24 | 0.807 | 0.958 | -0.01 | -0.40 | 0.690 | 0.972 |
| 18 | 0.00 | 0.05 | 0.958 | 0.958 | 0.00 | 0.13 | 0.896 | 0.972 |
| 19 | -0.05 | -1.96 | 0.050 | 0.958 | -0.03 | -1.35 | 0.178 | 0.972 |
| 20 | -0.03 | -1.43 | 0.154 | 0.958 | 0.00 | -0.21 | 0.836 | 0.972 |
| 21 | -0.02 | -1.03 | 0.305 | 0.958 | 0.03 | 1.33 | 0.183 | 0.972 |
| 22 | 0.00 | -0.07 | 0.944 | 0.958 | -0.04 | -1.60 | 0.110 | 0.972 |
| 23 | 0.01 | 0.61 | 0.545 | 0.958 | 0.00 | -0.17 | 0.863 | 0.972 |
| 25 | 0.00 | 0.16 | 0.871 | 0.958 | 0.03 | 1.28 | 0.200 | 0.972 |
| 26 | -0.04 | -1.63 | 0.104 | 0.958 | 0.01 | 0.47 | 0.638 | 0.972 |
| 27 | -0.06 | -2.41 | **0.016** | 0.566 | 0.03 | 1.36 | 0.175 | 0.972 |
| 28 | -0.02 | -0.88 | 0.378 | 0.958 | -0.02 | -0.93 | 0.352 | 0.972 |
| 29 | -0.01 | -0.52 | 0.600 | 0.958 | -0.04 | -1.76 | 0.079 | 0.972 |
| 30 | 0.00 | 0.18 | 0.859 | 0.958 | -0.04 | -1.60 | 0.109 | 0.972 |
| 31 | 0.01 | 0.23 | 0.819 | 0.958 | -0.09 | -3.66 | **<0.001** | **0.010** |
| 32 | -0.05 | -1.92 | 0.055 | 0.958 | 0.01 | 0.57 | 0.570 | 0.972 |
| 33 | 0.05 | 2.15 | **0.031** | 0.958 | 0.02 | 0.94 | 0.347 | 0.972 |
| 34 | -0.02 | -0.96 | 0.335 | 0.958 | 0.08 | 3.38 | **0.001** | **0.028** |

| | | | | | | | | |
|---|---|---|---|---|---|---|---|---|
| 35 | 0.07 | 2.82 | **0.005** | 0.180 | 0.01 | 0.48 | 0.629 | 0.972 |
| 36 | -0.02 | -0.89 | 0.373 | 0.958 | 0.00 | 0.07 | 0.943 | 0.972 |
| 37 | 0.04 | 1.60 | 0.109 | 0.958 | 0.05 | 2.17 | **0.030** | 0.972 |
| 38 | -0.02 | -0.95 | 0.344 | 0.958 | 0.00 | -0.05 | 0.960 | 0.972 |
| 39 | -0.01 | -0.49 | 0.625 | 0.958 | 0.02 | 1.03 | 0.301 | 0.972 |
| 40 | 0.02 | 0.90 | 0.367 | 0.958 | -0.03 | -1.28 | 0.200 | 0.972 |

The table shows effect sizes, t-values, and uncorrected (Unadj.) and corrected (Adj.) parametric p-values from the statistical analyses of the association between pregnancy-related dimensions and cortical components (CC)s, in the sensitivity analysis excluding variables with near-zero variance. Significant p-values are shown in bold.

**Table S7.** Associations between birth-related dimensions and cortical components from sensitivity analysis excluding variables with near-zero variance.

| Cortical Component | Low Birth Weight and Prematurity | | | | Newborn Birth Complications | | | |
|---|---|---|---|---|---|---|---|---|
| | Cohen´s D | t-value | p-value (unadj.) | p-value (adj.) | Cohen´s D | t-value | p-value (unadj.) | p-value (adj.) |
| 1 | -0.13 | -5.67 | **<0.001** | **<0.001** | 0.06 | 2.69 | **0.007** | 0.249 |
| 2 | -0.03 | -1.29 | 0.197 | 0.982 | 0.00 | -0.19 | 0.852 | 0.989 |
| 3 | 0.01 | 0.32 | 0.745 | 0.982 | -0.03 | -1.17 | 0.241 | 0.989 |
| 4 | -0.08 | -3.16 | **0.002** | 0.051 | 0.03 | 1.16 | 0.246 | 0.989 |
| 5 | -0.03 | -1.33 | 0.183 | 0.982 | 0.01 | 0.61 | 0.540 | 0.989 |
| 6 | -0.02 | -0.74 | 0.458 | 0.982 | 0.04 | 1.80 | 0.071 | 0.989 |
| 7 | -0.09 | -3.39 | **0.001** | **0.023** | 0.05 | 2.40 | **0.016** | 0.535 |
| 8 | 0.01 | 0.38 | 0.703 | 0.982 | -0.04 | -1.71 | 0.086 | 0.989 |
| 9 | 0.08 | 3.12 | **0.002** | 0.055 | -0.01 | -0.31 | 0.760 | 0.989 |
| 10 | -0.06 | -2.48 | **0.013** | 0.396 | 0.07 | 3.34 | **0.001** | **0.032** |
| 11 | 0.00 | -0.09 | 0.926 | 0.982 | 0.00 | 0.13 | 0.893 | 0.989 |
| 12 | 0.00 | -0.02 | 0.982 | 0.982 | 0.00 | 0.02 | 0.985 | 0.989 |
| 13 | 0.04 | 1.52 | 0.128 | 0.982 | 0.00 | 0.01 | 0.989 | 0.989 |
| 14 | -0.01 | -0.60 | 0.548 | 0.982 | -0.01 | -0.33 | 0.743 | 0.989 |
| 15 | -0.06 | -2.47 | **0.014** | 0.396 | 0.05 | **2.09** | **0.037** | 0.989 |
| 16 | 0.16 | 6.19 | **<0.001** | **<0.001** | -0.09 | -4.26 | **<0.001** | **0.001** |
| 17 | 0.02 | 0.57 | 0.570 | 0.982 | -0.02 | -0.75 | 0.453 | 0.989 |
| 18 | -0.09 | -3.63 | **<0.001** | **0.010** | 0.07 | 3.30 | **0.001** | **0.035** |
| 19 | -0.02 | -0.66 | 0.506 | 0.982 | -0.06 | -2.49 | **0.013** | 0.431 |
| 20 | -0.01 | -0.43 | 0.666 | 0.982 | 0.00 | 0.14 | 0.888 | 0.989 |
| 21 | -0.03 | -1.28 | 0.201 | 0.982 | 0.03 | 1.16 | 0.246 | 0.989 |
| 22 | 0.03 | 1.09 | 0.276 | 0.982 | -0.02 | -0.80 | 0.426 | 0.989 |
| 23 | -0.03 | -1.22 | 0.224 | 0.982 | -0.01 | -0.53 | 0.594 | 0.989 |
| 25 | 0.03 | 1.20 | 0.228 | 0.982 | -0.01 | -0.39 | 0.700 | 0.989 |
| 26 | -0.05 | -1.89 | 0.059 | 0.982 | 0.04 | 1.77 | 0.076 | 0.989 |
| 27 | -0.03 | -1.16 | 0.247 | 0.982 | 0.04 | 2.01 | **0.045** | 0.989 |
| 28 | -0.01 | -0.56 | 0.572 | 0.982 | 0.04 | 1.92 | 0.055 | 0.989 |
| 29 | 0.01 | 0.53 | 0.599 | 0.982 | -0.01 | -0.64 | 0.521 | 0.989 |
| 30 | -0.02 | -0.82 | 0.410 | 0.982 | -0.01 | -0.48 | 0.633 | 0.989 |
| 31 | 0.04 | 1.38 | 0.168 | 0.982 | -0.05 | -2.13 | **0.033** | 0.989 |
| 32 | -0.12 | -4.60 | **<0.001** | **<0.001** | 0.05 | 2.26 | **0.024** | 0.760 |
| 33 | 0.06 | 2.23 | **0.026** | 0.701 | -0.06 | -2.76 | **0.006** | 0.206 |

| | | | | | | | | |
|---|---|---|---|---|---|---|---|---|
| 34 | -0.06 | -2.24 | **0.025** | 0.701 | 0.02 | 0.85 | 0.398 | 0.989 |
| 35 | 0.04 | 1.39 | 0.165 | 0.982 | -0.03 | -1.45 | 0.148 | 0.989 |
| 36 | 0.05 | 1.79 | 0.073 | 0.982 | -0.03 | -1.48 | 0.139 | 0.989 |
| 37 | 0.14 | 5.92 | **<0.001** | **<0.001** | -0.02 | -0.77 | 0.438 | 0.989 |
| 38 | 0.02 | 0.93 | 0.353 | 0.982 | -0.03 | -1.34 | 0.182 | 0.989 |
| 39 | -0.14 | -5.34 | **<0.001** | **<0.001** | 0.05 | 2.11 | **0.035** | 0.989 |
| 40 | 0.01 | 0.41 | 0.681 | 0.982 | -0.03 | -1.16 | 0.245 | 0.989 |

The table shows effect sizes, t-values, and uncorrected (Unadj.) and corrected (Adj.) parametric p-values from the statistical analyses of the association between birth-related dimensions and cortical components (CC)s, in the sensitivity analysis excluding variables with near-zero variance. Significant p-values are shown in bold.

**Table S8.** Results of the Jarque-Bera Test for Normality

| | Maternal Pregnancy Complications | Maternal Substance Use | Low Birth Weight and Prematurity | Newborn Birth Complications |
|---|---|---|---|---|
| CC1 | 5272 **(<0.001)** | 5340 **(<0.001)** | 5418 **(<0.001)** | 5248 **(<0.001)** |
| CC2 | 7354 **(<0.001)** | 7468 **(<0.001)** | 7279 **(<0.001)** | 7374 **(<0.001)** |
| CC3 | 2750 **(<0.001)** | 2735 **(<0.001)** | 2751 **(<0.001)** | 2760 **(<0.001)** |
| CC4 | 29746 **(<0.001)** | 29690 **(<0.001)** | 29771 **(<0.001)** | 29787 **(<0.001)** |
| CC5 | 29153 **(<0.001)** | 29129 **(<0.001)** | 29099 **(<0.001)** | 29173 **(<0.001)** |
| CC6 | 22893 **(<0.001)** | 22890 **(<0.001)** | 22926 **(<0.001)** | 22905 **(<0.001)** |
| CC7 | 2051 **(<0.001)** | 2055 **(<0.001)** | 2061 **(<0.001)** | 2039 **(<0.001)** |
| CC8 | 410 **(<0.001)** | 410 **(<0.001)** | 412 **(<0.001)** | 410 **(<0.001)** |
| CC9 | 8308 **(<0.001)** | 8390 **(<0.001)** | 8289 **(<0.001)** | 8326 **(<0.001)** |
| CC10 | 6 (0.058) | 6 (0.053) | 5 (0.070) | 5 (0.078) |
| CC11 | 379 **(<0.001)** | 380 **(<0.001)** | 380 **(<0.001)** | 380 **(<0.001)** |
| CC12 | 66 **(<0.001)** | 67 **(<0.001)** | 67 **(<0.001)** | 67 **(<0.001)** |
| CC13 | 117 **(<0.001)** | 117 **(<0.001)** | 118 **(<0.001)** | 117 **(<0.001)** |
| CC14 | 6197 **(<0.001)** | 6180 **(<0.001)** | 6193 **(<0.001)** | 6191 **(<0.001)** |
| CC15 | 4717 **(<0.001)** | 4726 **(<0.001)** | 4622 **(<0.001)** | 4698 **(<0.001)** |
| CC16 | 574 **(<0.001)** | 579 **(<0.001)** | 612 **(<0.001)** | 584 **(<0.001)** |
| CC17 | 23 **(<0.001)** | 23 **(<0.001)** | 23 **(<0.001)** | 23 **(<0.001)** |
| CC18 | 1037 **(<0.001)** | 1038 **(<0.001)** | 1045 **(<0.001)** | 1035 **(<0.001)** |
| CC19 | 360 **(<0.001)** | 357 **(<0.001)** | 358 **(<0.001)** | 355 **(<0.001)** |
| CC20 | 3536 **(<0.001)** | 3570 **(<0.001)** | 3570 **(<0.001)** | 3571 **(<0.001)** |
| CC21 | 32 **(<0.001)** | 33 **(<0.001)** | 33 **(<0.001)** | 33 **(<0.001)** |
| CC22 | 120 **(<0.001)** | 123 **(<0.001)** | 121 **(<0.001)** | 121 **(<0.001)** |
| CC23 | 55 **(<0.001)** | 55 **(<0.001)** | 56 **(<0.001)** | 55 **(<0.001)** |
| CC25 | 38628 **(<0.001)** | 38639 **(<0.001)** | 38586 **(<0.001)** | 38684 **(<0.001)** |
| CC26 | 101 **(<0.001)** | 102 **(<0.001)** | 101 **(<0.001)** | 101 **(<0.001)** |
| CC27 | 61751 **(<0.001)** | 61457 **(<0.001)** | 61765 **(<0.001)** | 61697 **(<0.001)** |
| CC28 | 4021 **(<0.001)** | 4028 **(<0.001)** | 4031 **(<0.001)** | 4004 **(<0.001)** |
| CC29 | 194 **(<0.001)** | 201 **(<0.001)** | 202 **(<0.001)** | 202 **(<0.001)** |
| CC30 | 485 **(<0.001)** | 485 **(<0.001)** | 485 **(<0.001)** | 485 **(<0.001)** |
| CC31 | 593 **(<0.001)** | 593 **(<0.001)** | 593 **(<0.001)** | 587 **(<0.001)** |
| CC32 | 88 **(<0.001)** | 87 **(<0.001)** | 89 **(<0.001)** | 85 **(<0.001)** |
| CC33 | 74 **(<0.001)** | 74 **(<0.001)** | 74 **(<0.001)** | 76 **(<0.001)** |
| CC34 | 194 **(<0.001)** | 193 **(<0.001)** | 190 **(<0.001)** | 195 **(<0.001)** |
| CC35 | 96 **(<0.001)** | 94 **(<0.001)** | 95 **(<0.001)** | 93 **(<0.001)** |
| CC36 | 1139 **(<0.001)** | 1141 **(<0.001)** | 1141 **(<0.001)** | 1152 **(<0.001)** |
| CC37 | 1683 **(<0.001)** | 1678 **(<0.001)** | 1679 **(<0.001)** | 1701 **(<0.001)** |
| CC38 | 231 **(<0.001)** | 231 **(<0.001)** | 230 **(<0.001)** | 232 **(<0.001)** |
| CC39 | 73 **(<0.001)** | 73 **(<0.001)** | 74 **(<0.001)** | 74 **(<0.001)** |

| | | | | |
|---|---|---|---|---|
| CC40 | 103 **(<0.001)** | 103 **(<0.001)** | 103 **(<0.001)** | 104 **(<0.001)** |

Jarque-Bera statistic (p-value) from the Jarque-Bera test for normality of residuals from the linear mixed models. P-values in bold are significant at 0.05 threshold. The jarque.test from the moments package (version 0.14.1) in R was used.
